# Transcriptional, post-transcriptional, and post-translational mechanisms rewrite the tubulin code during cardiac hypertrophy and failure

**DOI:** 10.1101/2022.01.24.477567

**Authors:** Sai Aung Phyo, Keita Uchida, Christina Yingxian Chen, Matthew A. Caporizzo, Kenneth Bedi, Joanna Griffin, Kenneth Margulies, Benjamin L. Prosser

**Author notes:** **Correspondence:** Benjamin L. Prosser, Ph.D., Department of Physiology, Pennsylvania Muscle Institute, Cardiovascular Institute, University of Pennsylvania Perelman School of Medicine, Clinical Research Bldg Room 726, 415 Curie Blvd, Philadelphia PA 19104.

## Abstract

A proliferated and post-translationally modified microtubule network underlies cellular growth in cardiac hypertrophy and contributes to contractile dysfunction in heart failure. Yet how the heart achieves this modified network is poorly understood. Determining how the “tubulin code” – the permutations of tubulin isoforms and post-translational modifications - is rewritten upon cardiac stress may provide new targets to modulate cardiac remodeling. Further, while tubulin can autoregulate its own expression, it is unknown if autoregulation is operant in the heart or tuned in response to stress. Here we use heart failure patient samples and murine models of cardiac remodeling to interrogate transcriptional, autoregulatory, and post-translational mechanisms that contribute to microtubule network remodeling at different stages of heart disease. We find that autoregulation is operant across tubulin isoforms in the heart and leads to an apparent disconnect in tubulin mRNA and protein levels in heart failure. We also find that within 4 hours of a hypertrophic stimulus and prior to cardiac growth, microtubule detyrosination is rapidly induced to help stabilize the network. This occurs concomitant with rapid transcriptional and autoregulatory activation of specific tubulin isoforms and microtubule motors. Upon continued hypertrophic stimulation, there is an increase in post-translationally modified microtubule tracks and anterograde motors to support cardiac growth, while total tubulin content increases through progressive transcriptional and autoregulatory induction of tubulin isoforms. Our work provides a new model for how the tubulin code is rapidly rewritten to establish a proliferated, stable microtubule network that drives cardiac remodeling, and provides the first evidence of tunable tubulin autoregulation during pathological progression.

## Introduction

Heart Failure (HF) is a complex pathological condition in which cardiac performance fails to match systemic demand. HF is commonly preceded by an enlargement of the heart known as cardiac hypertrophy, which serves as a major risk factor for progression to HF. As such, understanding the molecular determinants of hypertrophy may reveal novel targets for HF prevention.

Microtubules are hollow tubes formed from the polymerization of α- and β-tubulin dimers that play essential roles in the structural support of cells, intracellular transport, and cell division. They exhibit stochastic growth and shrinkage and maintain a dynamic equilibrium between free and polymerized tubulin (S. Fig. 1A). Through their trafficking role, microtubules regulate cardiomyocyte electrical activity, mitochondrial dynamics, protein degradation and local translation, while also forming load-bearing structures that influence myocyte mechanics and mechano-signaling(Caporizzo et al. 2019).

During cardiac hypertrophy and heart failure, the microtubule network is significantly remodeled and acts as a double-edged sword. On one hand, a proliferated, stable microtubule network is essential for the development of cardiac hypertrophy in response to stressors such as adrenergic stimulation and hemodynamic overload(Sato et al. 1997; Fassett et al. 2009, 2019; Scarborough et al. 2021). Upon such hypertrophic stimuli, a dense microtubule network and the anterograde motor protein kinesin-1 coordinates the trafficking of mRNA and the translational machinery to control local synthesis and integration of nascent proteins(Scarborough et al. 2021). In the absence of microtubules, increased protein translation is decoupled from protein integration and the heart fails to grow(Scarborough et al. 2021), identifying an essential role of microtubule-based transport in adaptive cardiac growth.

Yet upon chronic stress, the densified microtubule network can also contribute to contractile dysfunction in HF(Tsutsui et al. 1999; Caporizzo et al. 2018; Chen et al. 2018). A collective body of research has established a causal link between aberrant microtubule network remodeling and impaired cardiac mechanics in HF. Tubulin mass, and consequently microtubule network density, is consistently increased in the myocardium of HF patients(Chen et al. 2018; Schuldt et al. 2020) and pressure-overloaded animals(Sato et al. 1997; Fassett et al. 2019), and its destabilization can improve dysfunctional cardiac mechanics(Tsutsui et al. 1993; Cheng et al. 2008; Chen et al. 2018; Caporizzo et al. 2020).

While the state of the microtubule network in advanced HF has been well-defined by recent studies(Chen et al. 2018; Schuldt et al. 2020), we know little about the drivers and temporal progression of changes to the microtubule network that occur during cardiac remodeling. A seemingly obvious mechanism to increase tubulin mass is transcriptional upregulation; yet when we examine published transcriptomic and proteomic data from HF samples, we observe a surprising but consistent inverse correlation between tubulin mRNA and protein levels across different causes of HF in multiple studies (Fig. 1A-B). This motivates a deeper examination between transcriptional and translation coupling of tubulin isoforms and other factors that could contribute to microtubule proliferation.

**Figure 1.**
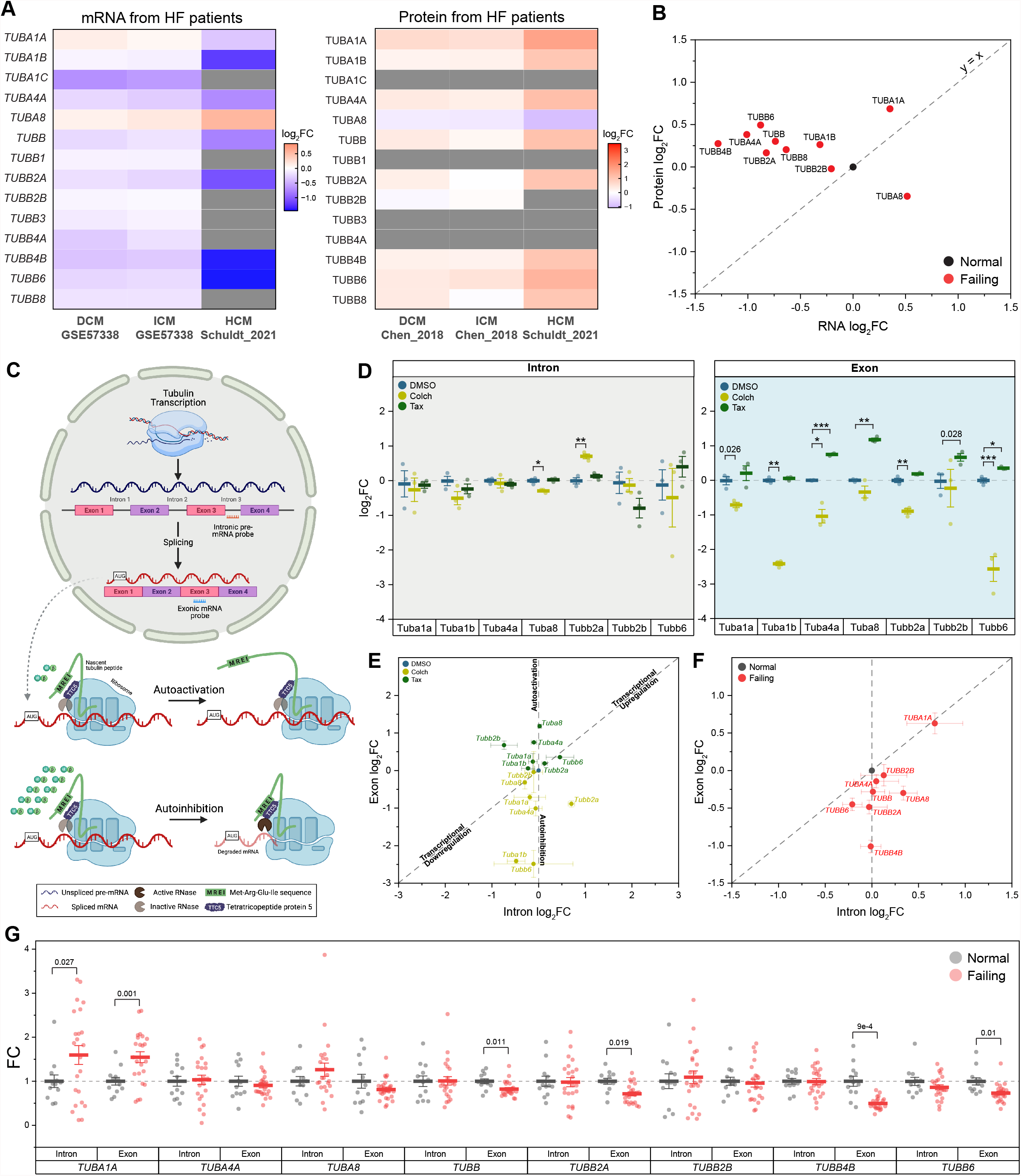
Tubulin autoregulation is operant in the heart and activated in heart failure. **(A)** Heatmaps of previously published mRNA (left) & protein (right) data of human αβ-tubulin isoforms during dilated (DCM)^8^, ischemic (ICM)^8^, & hypertrophic (HCM)^9^ cardiomyopathies. **(B)** Scatter-plot of log_2_fold-change of mRNA on x-axis and log_2_fold-change of protein on y-axis in heart failure; each data point represents an average log_2_fold-change value from DCM and ICM groups from (A), and y = x represents a proportionate change between mRNA and peptide. **(C)** Schematic of tubulin autoregulation: The introns of tubulin pre-mRNA are spliced out and the mRNA is fully translated in the absence of excess free tubulin; in the presence of excess free tubulin, the mRNA, but not the pre-mRNA, is degraded. **(D)** Relative log_2_fold-change of mRNA counts of intronic (left) and exonic (right) αβ-tubulin isoforms in isolated adult mouse cardiomyocytes after treatment with either depolymerizing (Colch) or polymerizing (Tax) agents (n=3); whiskers represent ± 1SEM, bolded line represents mean, * represents p-value from Welch-corrected two-tailed two-sample t-test on non-log data < 0.025 (Bonferroni-corrected for two comparisons), ** represents p < 0.01, and *** represents p < 0.001. **(E)** Scatter-plot of relative log_2_fold-change of intron on x-axis and exon on y-axis after Colch or Tax treatment in adult mouse cardiomyocytes; whiskers represent ± 1SEM. **(F)** Scatter-plot of relative log_2_fold-change of intron on x-axis and exon on y-axis in near-normal and failing patient heart samples; whiskers represent ± 1SEM. **(G)** Relative Fold-Change of mRNA counts of intronic and exonic αβ-tubulin isoforms in near-normal and failing patient heart samples; whiskers represent ± 1SEM, bolded line represents mean, and p-values are from Welch-corrected two-tailed two-sample t-test.

There are a multitude of α and β tubulin isoforms that arise from alternative tubulin genes; in humans, there are nine α and nine β -tubulin isoforms, and in mice, seven α and eight β isoforms (S. Fig. 1B). The abundance of tubulin transcripts can be controlled through autoregulation, a tubulin-specific mRNA rheostat in which an excess of free tubulin can activate a ribosomal RNase to degrade nascent tubulin transcripts (autoinhibition); conversely, if free tubulin levels are reduced, autoregulation is released (autoactivation) to promote tubulin synthesis and restore free tubulin content(Gasic and Mitchison 2019) (Fig. 1C). The extent to which tubulin isoforms are controlled through transcriptional or autoregulatory mechanisms has not been characterized, and autoregulation has not been examined in any capacity in the heart. Finally, any pathological relevance of autoregulation in cardiac or other tissues is largely unexplored.

The stabilization (i.e., protection from breakdown) of polymerized microtubules is another potentially important driver of the dense microtubule network observed in hypertrophy and HF. Microtubules are stabilized through association with microtubule-associated proteins (MAPs) and motors as well as through post-translational modifications (PTMs) of tubulin (S. Fig 1). Acetylation of polymerized α-tubulin produces long-lived and resilient microtubules that are resistant against repeated mechanical stresses(Kalebic et al. 2013) (Portran et al. 2017), while detyrosination - the removal of a tyrosine residue on the C-terminal tail of α-tubulin by vasohibins 1 & 2 (VASH1/2)(Aillaud et al. 2017; Nieuwenhuis et al. 2017) – stabilizes microtubules by modulating their interactions with depolymerizing effector proteins (Peris et al. 2009; Chen et al. 2021). The permutations of PTMs and tubulin isoforms is known as the “tubulin code” (S. Fig 1), which creates microtubule networks with distinct biochemical and mechanical properties. Altered detyrosination(Chen et al. 2018; Yu et al. 2021), acetylation(Swiatlowska et al. 2020), and MAP(Cheng et al. 2010; Li et al. 2018; Yu et al. 2021) binding are each implicated in pathological cardiac remodeling; yet how the tubulin code is rewritten during cardiac hypertrophy and HF remains largely unclear.

In this study, we interrogate changes to the tubulin code, MAPs, and motors at discrete stages of pathological cardiac remodeling. We find that surprisingly rapid and isoform-specific transcriptional induction and autoactivation of tubulin mRNA combine with post-translational detyrosination to drive microtubule stabilization and proliferation during early cardiac growth. We also find that in progressed heart failure, there is a switch to autoinhibition that reduces tubulin mRNA expression in the face of elevated tubulin protein content. This work identifies roles for autoregulation in rewriting the tubulin code during cardiac remodeling and may inform on approaches intended to modulate the course of hypertrophy and its progression to HF.

## Methods

### Human myocardial tissue

Procurement of human myocardial tissue was performed under protocols and ethical regulations approved by Institutional Review Boards at the University of Pennsylvania and the Gift-of-Life Donor Program (Pennsylvania, USA) and as described(Chen et al. 2018). In summary, failing human hearts were procured at the time of orthotropic heart transplantation at the Hospital of the University of Pennsylvania following informed consent from all participants. Non-failing hearts were obtained at the time of organ donation from cadaveric donors. In all cases, hearts were arrested in situ using ice-cold cardioplegia solution and transported on wet ice. Transmural myocardial samples were dissected from the mid left ventricular free wall below the papillary muscle and the samples were kept frozen at 80°C. Contractile parameters, including left ventricle ejection fraction, were determined by echocardiography in subjects. In this study, a total of 35 donor hearts were used. 12 donors were classified as near-normal non-failing (NF) without left-ventricular hypertrophy, and 23 donors were classified as heart failure with 12 hearts from hypertrophic cardiomyopathy patients and 11 hearts from dilated cardiomyopathy patients.

### Animal care

Animal care and procedures were approved and performed in accordance with the standards set forth by the University of Pennsylvania Institutional Animal Care and Use Committee (IACUC) and the Guide for the Care and Use of Laboratory Animals published by the US National Institutes of Health (NIH).

### Drug injection

Eight to twelve weeks old male C57/Bl6 mice were used throughout the study. On days 0 and 2, based on their body weights, mice were subcutaneously injected with either ascorbic acid (Ctrl, Sigma-Aldrich: A92902), 10mg/kg phenylephrine (PE, Sigma-Aldrich: P6126) prepared in Ctrl, or 5mg/kg isoproterenol (Iso, Sigma-Aldrich: I6504) prepared in Ctrl.

### Cardiac Tissue Harvest

Mice were put under general anesthesia using isoflurane and the hearts were surgically removed. Excised hearts were thoroughly washed in ice-cooled PBS and extra-cardiac tissues were removed. To properly measure heart weight (HW), residual blood from the chambers was removed by sandwiching the heart between Kimwipes and gently squeezing it. After HW measurement, atrial and right ventricular tissues were removed, the remaining septal and left-ventricular tissues were cut into five pieces of similar size and from similar locations of the heart. The weights of the individual pieces were recorded, frozen in liquid nitrogen, and stored at -80°C until further processing. Concurrent with tissue harvest, the tibia length (TL) of respective mouse was measured to calculate HW-over-TL (HW/TL).

### Exclusion criteria

During the study: for the 4-hour time point, there were 6 mice per treatment group for a total of 18 mice, and for the 4-day time point, there were 8 mice per treatment group for a total of 24 mice. As we aimed to study mice who underwent consistent cardiac hypertrophy, for the 4-day time point, we set exclusion criteria as the followings: (1) hearts whose HW/TL were beyond 2 standard-deviations (SD) of the population mean, and (2) experimental hearts whose classical hypertrophy response genes were not changed relative to that of the control hearts. After the removal of outliers, in the final study: for the 4-hour time point, there are 6 mice per treatment group for a total of 18 mice, and for the 4-day time point, there are 7 mice in Ctrl, 7 mice in PE, and 6 mice in Iso, for a total of 20 mice.

### Mouse cardiomyocyte isolation, culture, and drug treatment

Primary adult ventricular myocytes were isolated from 8- to 12-week-old C57/Bl6 mice using the protocol previously described(Prosser et al. 2011). Briefly, mice were put under general anesthesia using isoflurane and were injected peritoneally with heparin (∼1000 units/kg). The heart was excised and cannulated to a Langendorff apparatus for retrograde perfusion with enzymatic digestion solution at 37°C. Once digested, the heart was minced and triturated with glass pipettes. The isolated cardiomyocytes were centrifuged at 300 revolution per minute for 2 minutes. The supernatant containing debris was discarded and the isolated cells were resuspended in cardiomyocyte media containing Medium 199 (GIBCO: 11150-59) supplemented with 1x insulin-transferrin-selenium-X (GIBCO: 51500-56), 20mM HEPES pH 7.4, 0.1 mg/mL Primocin, and 25 μmol/L of cytochalasin D. Immediately following cell isolation, the cardiomyocytes were treated with either DMSO, 10μM colchicine, or 10μM taxol, and incubated at 37°C and 5% CO_2_ for 6 hours.

### Echocardiography

On day 4, transthoracic echocardiography was performed on mice, which were anesthetized using intraperitoneal injection of 0.01mL/gram body-weight of 2.5% Avertin, using Vevo2100 Ultrasound System (VisualSonics Inc., Toronto, Ontario, Canada). Fractional shortening, chamber dimensions, and ventricular wall-thickness were measured from short axis M-mode images at the mid-level view of the papillary muscle.

### Total protein lysate preparation

Frozen aliquoted cardiac tissue obtained from similar locations of the heart was pulverized finely using a liquid nitrogen-cooled mortar and pestle. 1x Radioimmunoprecipitation assay (RIPA) buffer (Cayman Chemical Company: 10010263) supplemented with 1x protease inhibitor cocktail (Cell Signaling Technology: 5872S) and 1:200 diluted endonuclease (Lucigen: OC7850K) was immediately added to the pulverized tissue at a constant ratio of 15μL/mg of tissue. The sample was then mechanically homogenized using a handheld homogenizer until visible chunks of tissues were dissociated. The sample was incubated for 10 minutes on ice to allow endonuclease to cleave DNA. After processing of all samples, the samples underwent two freeze-thaw cycles, after which, equal-volume of 5% SDS-10% glycerol boiling (SGB) buffer was added to each sample. The samples were vortexed thoroughly then heated to 100°C for 8 minutes. Residual undissolved cell debris were removed from the resulting samples by centrifugation at 8000g for 5 minutes at room temperature (22°C). The concentrations of the total protein were determined using Bicinchoninic acid (BCA) assay; all samples were diluted to 4μg/μL using RIPA:SGB buffer. The diluted total protein lysates were aliquoted and stored at - 80°C until further processing.

### Microtubule Fractionation

100mM PIPES-KOH pH 6.8, 1mM MgCl_2_, 1mM EGTA-KOH pH 7.7 (PME) buffer was prepared fresh and was supplemented with 1mM DTT, 1mM GTP (Sigma-Aldrich: G8877), and 1x protease inhibitor cocktail. 7 parts supplemented PME buffer was mixed with 3 parts glycerol; the final PME-30G buffer was kept incubated in a 37°C water bath. Frozen aliquoted cardiac tissue obtained from similar locations of the heart was pulverized crudely using a liquid nitrogen-cooled mortar and pestle. Immediately following pulverization, warmed PME-30G buffer was added at a constant ratio of 20μL/mg of tissue. The sample was then mechanically homogenized using the handheld homogenizer until visible chunks of tissues were dissociated and was set aside at 22°C until all samples were processed. All processed samples were then centrifuged at 16000g for 15 minutes at 30°C; the supernatants were transferred into fresh tubes and were saved as free tubulin (Free) fractions. 10μL of 1 part RIPA and 1 part SGB (RIPA:SGB) buffer was added to the pellet obtained from 1mg of tissue and the sample was homogenized using the handheld homogenizer. After processing of all samples, the samples were heated to 100°C for 8 minutes, cooled on ice, and centrifuged at 8000g for 5 minutes at 22°C; the supernatants were transferred into fresh tubes and were saved as polymerized tubulin (Poly) fractions. The concentrations of the Poly fractions were determined using BCA assay. The Poly fractions were diluted to 4μg/μL using RIPA:SGB buffer; the respective Free fraction was diluted with PME-30G buffer using twice the volume needed to dilute the Poly fraction. The final diluted Free and Poly fractions were aliquoted and stored at -80°C until further processing.

### Sample preparations and Western blot (WB) analysis

To quantify the relative abundance of specific proteins of interest in the total protein lysate, aliquoted diluted total protein lysate samples were thawed at 22°C. 1 part 4x loading buffer (125mM Tris-HCl pH 6.8, 35% v/v glycerol, 0.2% w/v Orange G) freshly supplemented with 10% v/v β-mercepthoethanol (BME) was mixed with 3 parts total protein lysate to get final concentrations of 1x loading buffer with 2.5% BME, and 3μg/μL of total protein. The final samples were heated to 100°C for 8 minutes. The heated samples were cooled to 22°C, centrifuged briefly, vortexed thoroughly, and loaded 5μL/sample onto precast protein gels (Bio-Rad: 5671085).

To quantify the relative abundances of the Free and Poly fractions, aliquoted diluted Free and Poly fractions were thawed at 22°C. For the Free fractions, 2x loading buffer (62.5mM Tris-HCL pH 6.8, 5% v/v SDS, 0% glycerol, 0.1% w/v Orange G) freshly supplemented with 5% v/v BME was used, whereas, for the Poly fractions, 4x loading buffer freshly supplemented with 10% v/v BME was used; to prepare the final samples, the respective loading buffers were diluted to 1x using the Free and Poly fractions. The final samples were heated to 100°C for 8 minutes. The heated samples were cooled to 22°C, centrifuged briefly, vortexed thoroughly, and loaded 5μL/Poly fraction and 10μL/Free fraction onto precast protein gels.

Protein gel electrophoresis was carried out under constant voltage of 135V for the Midi gels for 1 hour. The resolved proteins were transferred onto a nitrocellulose membrane using the Turbo Transfer System (Bio-Rad) under recommended conditions. The post-transferred membrane was blocked in blocking buffer (LI-COR Biosciences: 927-60003) for at least 1 hour at 22°C (or overnight at 4°C). The blocked membrane was incubated overnight at 4°C with primary antibodies diluted in 1x Tris buffered saline with Tween-20 (TBST, Cell Signaling Technology: 9997S). The membrane was washed twice using TBST, and incubated for 1 hour at 22°C with secondary antibodies diluted in blocking buffer. The final immunoblotted membrane was washed twice using TBST and was imaged using the Odyssey Western Blot Imaging System (LI-COR Biosciences).

### WB data analysis

The WB data was analyzed using Image Studio Lite (LI-COR Biosciences). The signal intensity of an individual band was obtained by drawing a rectangular block encompassing the entire band. The background was thresholded using the parameters: median, border width = 3, Top/Bottom. 2 technical replicates (n) per sample, and 6 biological replicates (N) per treatment for 4-hour time point and 8 biological replicates per treatment for 5-day time point were used in the analysis. GAPDH intensity was used as a loading control. A mean value of the Ctrls that were run on the same blot was used to normalize the data and to calculate the relative fold-changes over the Ctrl. Statistical analyses were performed as described below.

### Primary and secondary antibodies

| Target | Vendor | Host Species | Clonal | Product No. | Concentration used in WB (µg/mL) |
| --- | --- | --- | --- | --- | --- |
| Total α-tubulin | Abcam | Mouse | Mono | ab7291 | 0.33 |
| Total α-tubulin | Abcam | Rabbit | Poly | ab4074 | 0.5 |
| Total β-tubulin | Abcam | Rabbit | Poly | ab6046 | 0.67 |
| Acetylated α-tubulin | Abcam | Mouse | Mono | ab24610 | 1 |
| Detyrosinated α-tubulin | Abcam | Rabbit | Poly | ab48389 | 1 |
| Polyglutamylated α-tubulin | Adipogen | Mouse | Mono | 50-436-394 | 2 |
| Δ2 α-tubulin | Moutin Lab | Rabbit | Poly |  | 1:5000 |
| Polyglycylated tubulin | EMD Millipore | Mouse | Mono | MABS276 | 2 |
| GAPDH | GenScript | Mouse | Mono | A01622-40 | 0.5 |
| H3 | Abcam | Mouse | Mono | ab24834 | 0.33 |
| Anti-mouse | LI-COR | Donkey | Poly | 925-32212 | 0.1 |
| Anti-rabbit | LI-COR | Donkey | Poly | 925-68073 | 0.1 |

### Mass spectrometry (MS) sample preparation

To quantify the relative changes of multiple proteins of interest in the total protein lysate, aliquoted diluted total protein lysate samples were thawed at 22°C. 1 part 4x loading buffer freshly supplemented with 10% v/v BME was mixed with 3 parts total protein lysate. The final samples were heated to 100°C for 8 minutes. The heated samples were cooled to 22°C, centrifuged briefly, vortexed thoroughly, and loaded 50μL/sample onto precast protein gels (Bio-Rad: 4561034). Protein gel electrophoresis was carried out under constant voltage of 110V for the Mini gels for 1.5 hours. The resolved protein gel was stained with Coomassie blue (Bio-Rad: 1610435) using the provided protocol. After the destaining of the gel, the 50kDa bands were carefully excised and stored in deionized water at 4°C until further processing.

The gel bands were destained with 100mM ammonium bicarbonate/acetonitrile (50:50). The bands were reduced in 10mM dithiothreitol/100mM Ammonium bicarbonate for over 60 minutes at 52°C; the bands were then alkylated with 50mM iodoacetamide/100mM ammonium bicarbonate at 22°C for 1 hour in the dark. The proteins in the gel bands were digested with enzymes while incubating overnight at 37°C; different enzymes such as trypsin, Chymotrypsin, and Glu-C were used according to protein sequences. The supernatants were transferred and kept in fresh tubes. Additional peptides were extracted from the gel by adding 50% acetonitrile/1% TFA and incubated for 10 minutes on a shaker. The supernatants were combined and dried. The dried samples were reconstituted using 0.1% formic acid for MS analysis.

### MS analysis using Nano-LC-MS/MS

Peptides were analyzed on a Q-Exactive HF (Thermo Fisher Scientific) attached to an Ultimate 3000 rslcnano system (Thermo Fisher Scientific) at 400 nL/min. Peptides were eluted with a 55 minutes gradient from 5% to 32% ACN (25 minutes) and 90% ACN over 5 minutes in 0.1% formic acid. Data-dependent acquisition mode with a dynamic exclusion of 45 seconds was enabled. One full MS scan was collected with a scan range of 350 to 1200 *m*/*z*, resolution of 70 K, maximum injection time of 50 milliseconds, and AGC of 1 × 10^6^. Then, a series of MS2 scans were acquired for the most abundant ions from the MS1 scan (top 12). Ions were filtered with charges 2–4. An isolation window of 2 *m*/*z* was used with quadruple isolation mode. Ions were fragmented using higher-energy collisional dissociation (HCD) with a collision energy of 27%. Orbitrap detection was used with a scan range of 140 to 2000 *m*/*z*, resolution of 30 K, maximum injection time of 54 milliseconds, and AGC of 50,000.

### MS data analysis

Proteome Discoverer (Thermo Fisher Scientific, version 2.4) was used to process the raw spectra. Default search parameters were used, including precursor mass tolerance of 10 ppm, fragment mass tolerance of 0.02 Da, enzymes specific cleavage, and up to 2 mis-cleavage. Carbamidomethyl [C] was set as a fixed modification, while Oxidation [M] and Acetylation [N-terminal and K] were set as variable modifications. The target-decoy approach was used to filter the search results, in which the false discovery rate was less than 1% at the peptide and protein levels. For measuring the relative protein abundances, all the chromatographic data were aligned and normalized to peptide groups and protein abundances, missing values were imputed and scaled. Since the different tubulin isoforms share multiple homologies between one another, only unique peptides that are unambiguous to each isoform was used to calculate protein abundance. The unique peptide acquired for the analyzed isoforms range from 1-13 and the full suite of peptide and protein groups used in the analysis can be found in the public proteomic repository as outlined in the data availability statement. Statistical analyses were performed on the calculated protein abundances as described below.

### Total RNA extraction

Frozen aliquoted cardiac tissue obtained from similar locations of the heart was pulverized finely using a liquid nitrogen-cooled mortar and pestle. 500μL of ice-cooled RNAzol (Molecular Research Center: RN 190) was added to the pulverized tissue and immediately homogenized using the handheld homogenizer until visible chunks of tissues were dissociated. 200μL of molecular grade water was added to the sample; the sample was vortexed and incubated for 15 minutes at 22°C. After processing of all samples, the samples were then centrifuged at 12000g for 15 minutes at 22°C. 550μL of the clear supernatant was carefully removed and transferred into a fresh tube. 550μL of isopropanol was then added to the supernatant, vortexed, and incubated for 10 minutes at 22°C. The samples were centrifuged at 16000g for 10 minutes at 22°C, and the resulting supernatants were discarded. The visible RNA pellets were washed in 75% ethanol in molecular grade water three times. The undried RNA pellets were resuspended in 30μL of RNase free water. The total RNA concentrations, and 260/230 and 260/280 ratios were determined using NanoDrop ND-1000 Spectrophotometer (NanoDrop Technologies). The RNA samples were stored at -80°C until further analysis.

### NanoString nCounter analysis

Total RNAs from 37 samples were analyzed. The concentration of the total RNA was reassessed using NanoDrop spectrophotometer. The quality of the total RNA was assessed using the Agilent 4200 TapeStation (Agilent Technologies). Only samples that were pure as defined by OD 260/280 and 260/230 ratios > 1.8, and integrity RIN value > 8.0 were used in the study. 100ng of total RNA per sample for tubulin and hypertrophy panels or 200ng of total RNA per sample for tubulin autoregulation panel was used for the subsequent step. Hybridization between the target mRNA and reporter-capture probe pairs was performed for 18 hours at 65°C using Mastercycler Pro S Thermal Cycler (Eppendorf) according to the manufacturer’s protocol. Post-hybridization processing was carried out on a fully automated nCounter Prep Station (NanoString Technologies) liquid-handling robotic device using the High Sensitivity setting. For image acquisition and data processing, the probe/target complexes were immobilized on the nCounter cartridge that was then placed in the nCounter Digital Analyzer (NanoString Technologies) as per the manufacturer’s protocol with FOV set to 555. The expression level of a gene was measured by counting the number of times the probe with a unique barcode, which was targeted against that gene, was detected. The barcode counts were then tabulated in a comma-separated value (.csv) format.

### NanoString nCounter data and statistical analysis

The raw digital counts of expressions were exported into nSolver Analysis software (NanoString, version 4.0) for downstream analysis. The data was analyzed in nSolver using the Nanostring Analysis and Advanced Analysis software packages. The background of the data was thresholded using the geometric means after removing negative control values that are three-times higher than the rest. The data was then normalized using the geometric means of the positive controls, after removing “F” if the value is too close to background, and the three housekeeping genes (Gapdh, Rpl4, Tbp). Without removing low count values, the Bonferroni-corrected differentially expressed gene (DEG) analysis of the normalized data was computed using Treatment as covariates. For tubulin autoregulation panel, raw counts were exported, and statistical analyses were carried as outlined below.

### Statistical Analysis

Graphing and statistical analyses were performed using OrginPro 2019 software (OriginLabs). First, the normality of the data was determined using the <u>Shapiro-Wilk test</u>. For comparison of data distributions whose normality cannot be rejected at 0.05 level, the calculated probability of the means (p) between the control and the experimental group was calculated using the two-tailed two-sample Welch-corrected student’s t-test. For comparison of data distributions whose normality is rejected at 0.05 level, the p-value between the control and the experimental group was calculated using the two-tailed two-sample Kolmogorov-Smirnov test. For significance level, we used the Bonferroni-corrected significance cut-off of p < 0.025 denoted by * ; ** represents p < 0.01 and *** represents p < 0.001. P-values to two significant figures were reported for 0.05 < p < 0.025. For all bar graphs, the bar represents mean and the whisker represents + 1 SEM. For all box plots, the bolded line represents mean, and the whiskers represent ± 1 standard error of mean (SEM).

## Results

### Tubulin autoregulation is operant in the heart and induced in heart failure

Despite the importance of microtubule proliferation in cardiac pathology, any role of tubulin autoregulation has not been examined. We utilized a previously established strategy to test for autoregulation by measuring the relative abundances of pre-spliced (i.e. intron-containing) and spliced (i.e. those without introns) tubulin mRNAs(Gasic et al. 2019). Using this approach, one can detect transcriptional regulation of a target through correlated changes in intronic and exonic mRNA levels (y = x in Fig. 1E, for example), whereas post-transcriptional autoregulation would only affect exonic mRNA (shift along the y-axis of Fig. 1E). To study autoregulation in an isoform-specific fashion, we designed NanoString nCounter probes for direct and unique detection of either intronic or exonic regions of individual tubulin isoforms. To determine if autoregulation is operant in heart muscle cells, we treated isolated mouse cardiomyocytes for 6 hours with colchicine, a microtubule depolymerizing agent predicted to trigger autoinhibition (decrease in only exonic species) by increasing free tubulin, or taxol, a microtubule polymerizing agent predicted to trigger autoactivation (increase in only exonic species) by shifting free tubulin into the polymerized pool. Consistently, depolymerization significantly reduced the amount of exonic but not intronic mRNA across most tubulin isoforms, while polymerization increased the amount of exonic tubulin mRNA (Fig. 1D-E). This data serves as the first demonstration that autoregulation is operant in the cardiomyocyte and that it regulates the majority of tubulin isoforms.

Next, we tested whether tubulin autoregulation can partially explain the discrepancy in mRNA and protein levels observed in HF. To this end, we designed a separate Nanostring probe set against introns and exons of human tubulin isoforms and probed RNA extracted from 35 cardiac samples from 12 non-failing donors or 23 patients with advanced heart failure. Figure 1F-G shows the relative intronic and exonic abundances for all tubulin isoforms that could be readily detected at the intronic, exonic and protein level (Chen et al. 2018)(Figure 1A-B). In failing hearts, the majority of tubulin isoforms showed reduced exonic relative to intronic levels, indicative of active autoinhibition across most isoforms. An exception is *TUBA1A*, the only isoform that demonstrated significant transcriptional induction; consistently *TUBA1A* is also being the only isoform to show both increased and correlated mRNA and protein levels in this HF population (Figure 1A-B). Of additional note, *TUBB4B* is by far the most abundant α-tubulin isoform expressed in the heart, and it exhibits robust autoinhibition in HF, yet maintains increased protein abundance. Taken together this data indicates that in HF, elevated tubulin protein triggers persistent autoinhibition of tubulin mRNA. The maintained elevation in tubulin protein may be explained by significantly increased tubulin stability/lifetime.

However, there remains no explanation as to how the heart achieved the increased tubulin protein in the first place or whether autoregulation plays any role in the establishment of the increased tubulin mass observed in pathological cardiac remodeling. To better understand this, we employed mice models of cardiac hypertrophy that allows us to explore the early roles of tubulin transcription, autoregulation, and stability.

### Acute adrenergic agonism induces anatomic and transcriptional cardiac remodeling

To determine how the microtubule network remodels during the development of cardiac hypertrophy, we characterized the myocardial cytoskeleton at two time points in two mouse models of adrenergic agonist-induced hypertrophy(Scarborough et al. 2021) (Fig. 2A). A 4-hour post-injection time point was chosen to capture a stage when hearts were exposed to hypertrophic stimuli but have not yet hypertrophied, i.e. pre-hypertrophy, and a 4-day post-injection time point was chosen to capture a stage when hearts had demonstrably hypertrophied. As expected, no change in heart-weight-to-tibia-length (HW/TL) was observed 4 hours after injection of either phenylephrine (PE) or isoproterenol (Iso) compared to vehicle control (Ctrl) (Fig. 2B, left). When mice were given a second injection on day 2 and hearts were collected on day 4, we observed a consistent cardiac hypertrophy with both PE and Iso (Fig. 2B, right). To assess left-ventricular remodeling and function, we performed echocardiography on the 4-day hypertrophy animals. We observed consistent evidence of concentric hypertrophy upon both PE and Iso treatment, with elevated left-ventricular (LV) mass and increased wall and septal thickness (Fig. 2C-D). Neither group exhibited evidence of decompensation toward HF, with no evidence of ventricular dilation, or depressed contractility, indicating a compensated, concentric hypertrophy in response to acute adrenergic agonism.

**Figure 2.**
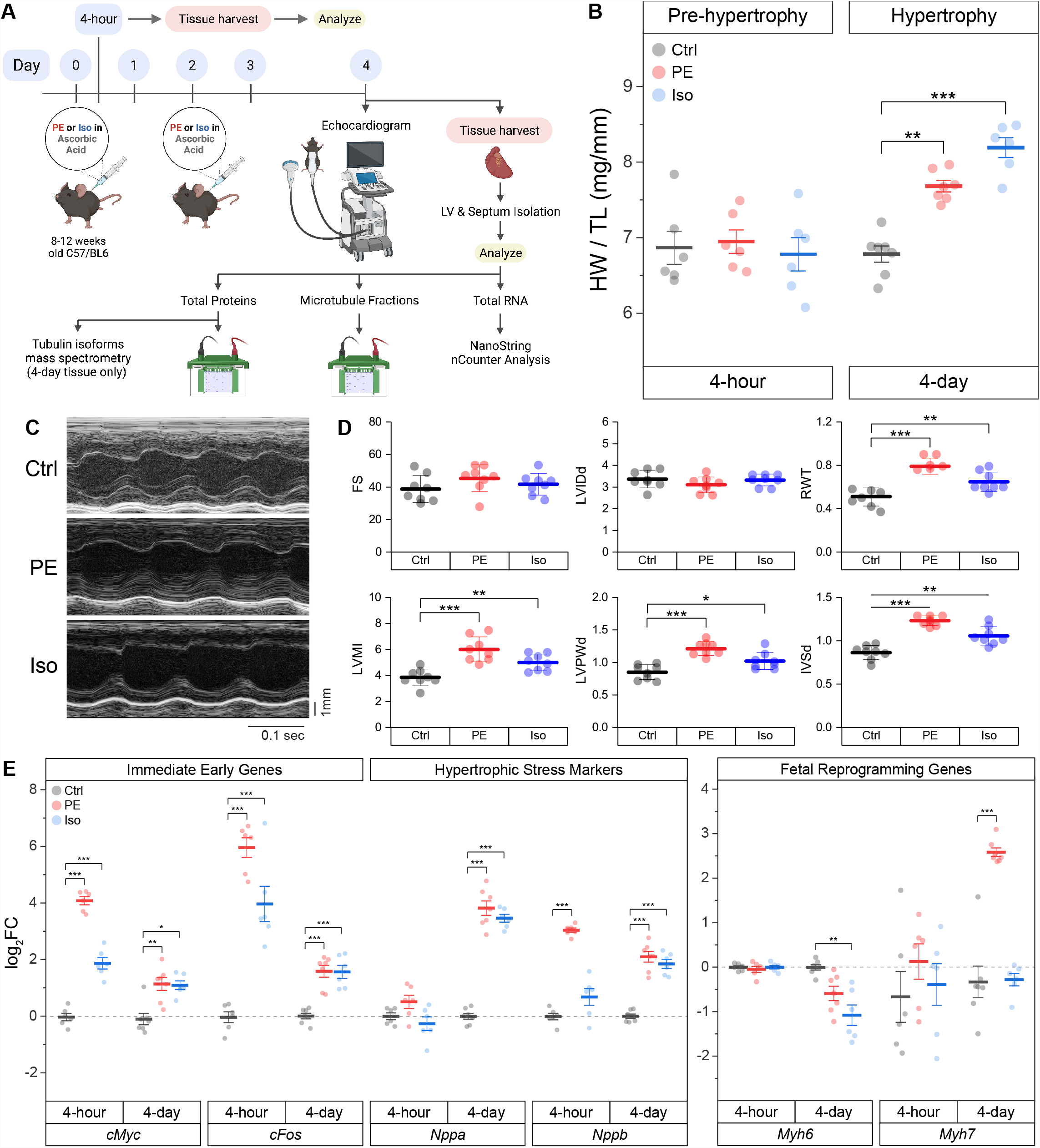
Acute α- or β-adrenergic stimulation induces cardiac hypertrophy. **(A)** Graphical scheme of the experimental plan. **(B)** Heart-weight / Tibia length (HW/TL) data of mice after 4-hour (pre-hypertrophy) (n=6) or 4-day (hypertrophy) (n = Ctrl:7, PE:7, Iso:6) following 10mg/kg/injection of phenylephrine (PE) or 5mg/kg/injection of isoproterenol (Iso). **(C)** Representative echocardiographic M-mode images of 4-day mice hearts. **(D)** Quantification of relevant echocardiographic parameters: FS = Fractional Shortening, LVIDd = Left-Ventricular Internal Diameter at end diastole, RWT = Relative Wall Thickness, LVMI = Left-Ventricular Mass Index, LVPWd = Left-Ventricular Posterior Wall thickness at end diastole, IVSd = InterVentricular Spetal thickness at end diastole (n=8). **(E)** Relative log_2_fold-change of nCounter mRNA counts of Immediate Early Genes (IEGs), hypertrophic stress markers, and genes of fetal reprogramming (n = 4h: 6, 4d: Ctrl:7, PE:7, Iso:6). For all box plots, whiskers represent ± 1SEM and bolded-lines represent mean. For (B) and (D), * represents p-value from Welch-corrected two-tailed two-sample student’s t-test < 0.025, ** represents p < 0.01, and *** represents p < 0.001. For (E), * represents Bonferroni adjusted (for 45 genes) p-value < 0.025 (Bonferroni-corrected for two comparisons), ** represents adj_p < 0.01, and *** represents adj_p < 0.001 (see Methods for more statistical details).

We further validated our models using NanoString nCounter to assess transcriptional markers of cardiac remodeling in the hearts of PE and Iso treated mice. Using direct RNA counting of 42 transcripts sorted into immediate early genes (IEGs), hypertrophy-related genes, and fibrosis-related genes, we analyzed differentially expressed genes (DEGs) in the septa and LV of our time-matched control and experimental groups. We hypothesized that IEGs would be upregulated after 4-hour of adrenergic stimulation, followed by upregulation of canonical markers of hypertrophic remodeling after 4 days. Consistent with this hypothesis, we observed robust upregulation of the canonical IEGs – *cMyc* and *cFos* – in both PE and Iso treated mice at 4-hour (Fig. 2E & S. Fig. 2), followed by induction of stress markers – *Nppa* and *Nppb* – at 4-day, along with markers of fetal reprogramming including myosin isoform switching (reduced *Myh6:Myh7* ratio).

The full complement of DEGs in each experimental group at the 4-hour and 4-day time points are depicted in S. Fig. 2. At both 4-hour and 4-day after adrenergic agonism, we observed upregulation of hypertrophy-related gene *Fhl1*(Friedrich et al. 2012), and fibrosis-related genes *Ctgf* (Hayata et al. 2008) & *Vcan*(Vistnes et al. 2014). Additional potentially relevant DEGs included the upregulation of *Col4a1*(Steffensen and Rasmussen 2018) and *Timp1*(Barton et al. 2003), and the downregulation of *Agrn*(Bassat et al. 2017; Baehr et al. 2020), among others.

### The microtubule network is rapidly detyrosinated upon hypertrophic stimulation

Having validated the 4-hour and 4-day models, we examined microtubule network remodeling in these contexts. We first determined whether the total αβ-tubulin content and free vs. polymerized tubulin pools are altered in the pre-hypertrophic state, i.e. 4-hour. We observed no significant differences in these metrics of total tubulin content or fractionation at this early time point (Fig. 3A-C).

**Figure 3.**
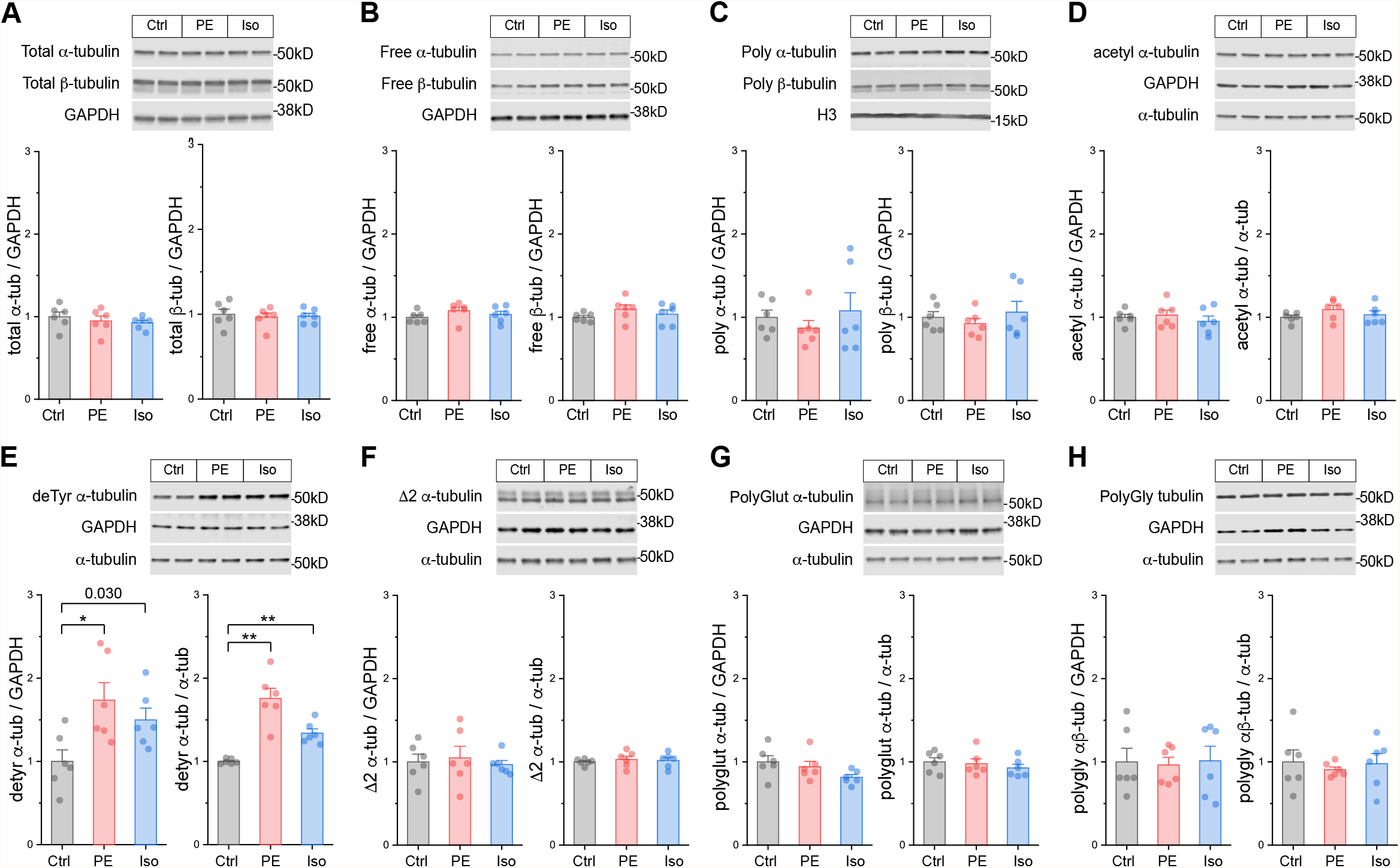
The microtubule network is rapidly detyrosinated upon hypertrophic stimulation. Representative immunblots and relative fold-change of α-tubulin **(left)** and β-tubulin **(right)** in **(A)** total proteins, **(B)** free or **(C)** polymerized -tubulin fractions (n=6). Representative immunoblots with technical duplicate lanes and relative fold-change using GAPDH **(left)** or α-tubulin **(right)** as loading controls for **(D)** acetylated α-tubulin, **(E)** detyrosinated α-tubulin, **(F)** Δ2 α-tubulin, **(G)** polyglutamylated α-tubulin, & **(H)** polyglycylated pan-tubulin (n=6). For all bar plots, whiskers represent + 1SEM and bar represents mean. For all graphs, * represents p-value from Welch-corrected two-tailed two-sample t-test < 0.025 (Bonferroni-corrected for two comparisons), ** represents p < 0.01, and *** represents p < 0.001.

We next determined whether tubulin is rapidly post-translationally modified upon hypertrophic stimulation. We immunoblotted for the five best-studied PTMs using validated antibodies: acetylation, detyrosination, polyglutamylation, polyglycylation, and Δ2 tubulin. Polyglutamylation, polyglycylation, and Δ2 are well characterized in cilia, flagella, and the brain(Paturle-Lafanechère et al. 1994; Aillaud et al. 2016), but they have not been studied in the heart. Detyrosination and acetylation, which occur predominantly on polymerized microtubules, are common markers of stable, long-lived microtubules, and of microtubule damage and repair-stabilization processes(Portran et al. 2017; Xu et al. 2017) respectively.

At the 4-hour time point, we did not observe any significant differences in either the absolute (PTM/GAPDH) or the relative (PTM/α-tubulin) amounts of acetylation, polyglutamylation, polyglycylation, or Δ2 tubulin (Fig. 3D, F-H). Surprisingly, we did observe robust induction of the absolute and relative amounts of detyrosination in both PE and Iso treated groups (Fig. 3E). These data suggest that within 4 hours of hypertrophic stimulation, prior to other overt changes in tubulin mass, microtubules are rapidly detyrosinated, which could be an early driver of microtubule stabilization.

### Post-translationally modified microtubules proliferate during the establishment of cardiac hypertrophy

We next characterized microtubule network remodeling at day 4, concurrent with cardiac hypertrophy. We probed the 3 tubulin pools and immunoblotted for α-tubulin, β-tubulin, acetylation, detyrosination, polyglutamylation, polyglycylation, Δ2 as described above.

At this stage we observed increased free, polymerized, and total αβ-tubulin protein in the hearts of PE and Iso-treated mice (Fig. 4A-C). In the PE group, the ratio of free:polymerized α-tubulin decreased (S. Fig. 3D), consistent with enhanced microtubule stability. In PE-treated mice, we observed increases in the absolute amounts of acetylation, polyglutamylation, and polyglycylation, and in the absolute and relative amounts of detyrosination. Iso treated mice showed a similar trend for each PTM, but of reduced magnitude and greater variability (Fig. 4C-H). Taken together, these data indicate that during cardiac hypertrophy tubulin content increases, the polymerized network densifies, and there is a proportionally increased abundance of post-translationally modified microtubules with a modest enrichment of detyrosination.

**Figure 4.**
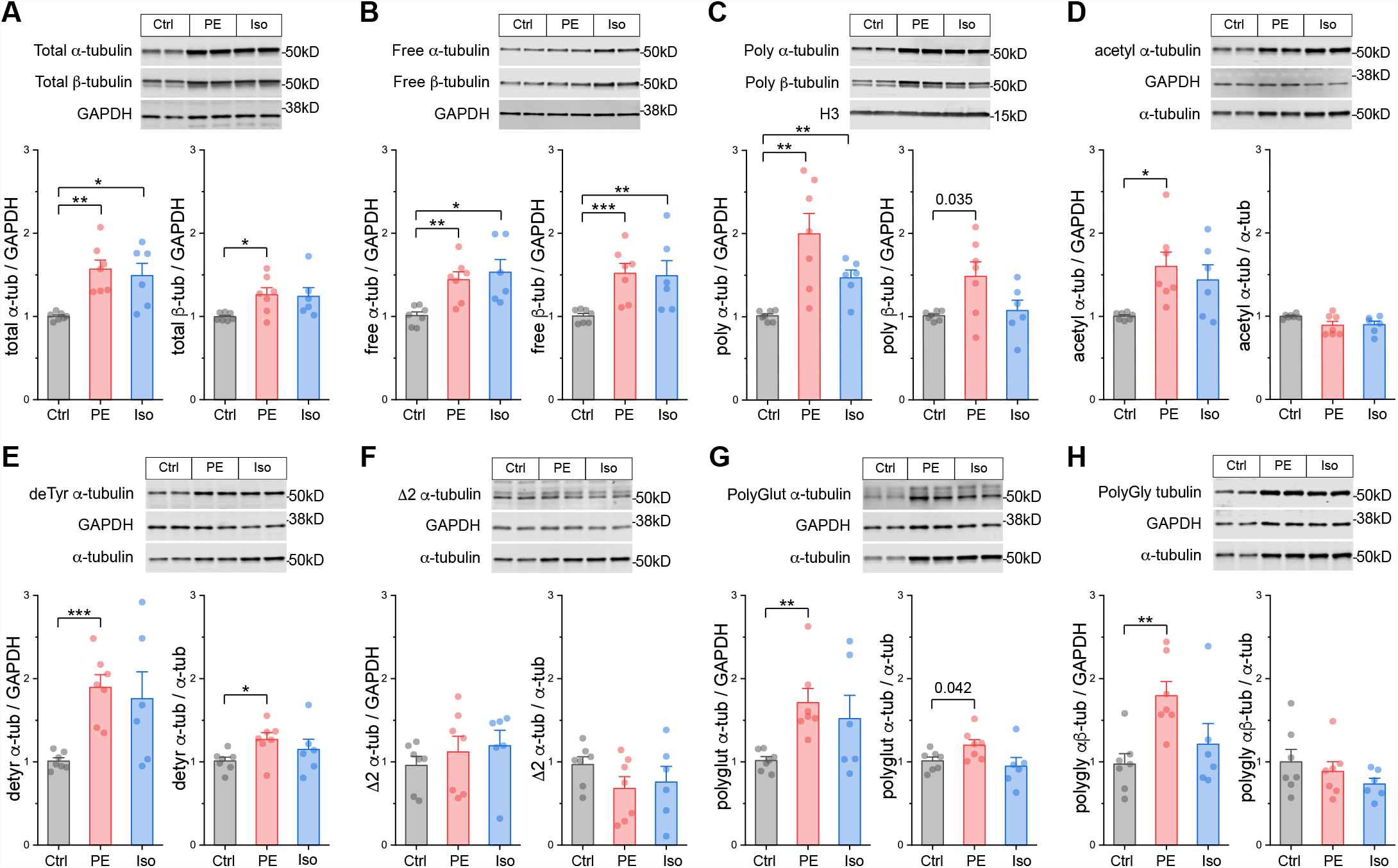
Total tubulin content increases & the microtubule network densifies during hypertrophy. Representative immunblots and relative fold-change of α-tubulin **(left)** and β-tubulin **(right)** in the **(A)** total proteins, **(B)** free, or **(C)** polymerized -tubulin fractions (n = Ctrl:7, PE:7, Iso:6). Representative immuno-blots with technical duplicate lanes and relative fold-change using GAPDH **(left)** or α-tubulin **(right)** as loading controls for **(D)** acetylated α-tubulin, **(E)** detyrosinated α-tubulin, **(F)** Δ2 α-tubulin, **(G)** polyglutamylated α-tubulin, & **(H)** polyglycylated pan-tubulin (n = Ctrl:7, PE:7, Iso:6). For all bar plots, whiskers represent + 1SEM and bar represents mean; * represents p-value from Welch-corrected two-tailed two-sample t-test < 0.025 (Bonferroni-corrected for two comparisons), ** represents p < 0.01, and *** represents p < 0.001.

We next sought to determine how specific tubulin isoforms contribute to the increase in tubulin content observed at 4-day. To this end, we utilized mass spectrometric (MS) analysis of the total tubulin pool. We observed that the predominant α- and β-tubulin isoforms of murine LV were Tuba1a and Tubb4b, respectively (Fig. 5A). Each of these predominant isoforms were modestly increased upon PE and Iso treatment. We also determined the relative changes of all detectable tubulin isoforms and observed significant increases in Tuba1a, Tuba1c, Tubb2a, Tubb2b, Tubb3, Tubb5, and Tubb6 (Fig. 5A-B). Of note, Tuba4a – the only tubulin isoform that is synthesized in its detyrosinated form – was clearly not increased upon hypertrophic stimulation. This indicates that the early increases in detyrosination are not due to increased synthesis of Tuba4a, and instead likely due to altered activity of the enzymes of the tyrosination cycle. Tubb6 exhibited the highest degree of upregulation with a ∼4-fold increase upon PE treatment; this is notable as Tubb6 induction has been causally implicated in microtubule network reorganization in Duchenne Muscular Dystrophy(Randazzo et al. 2019). Despite significant upregulation of multiple low abundance isoforms, the overall composition of the total tubulin pool is largely conserved at this stage of hypertrophic remodeling.

**Figure 5.**
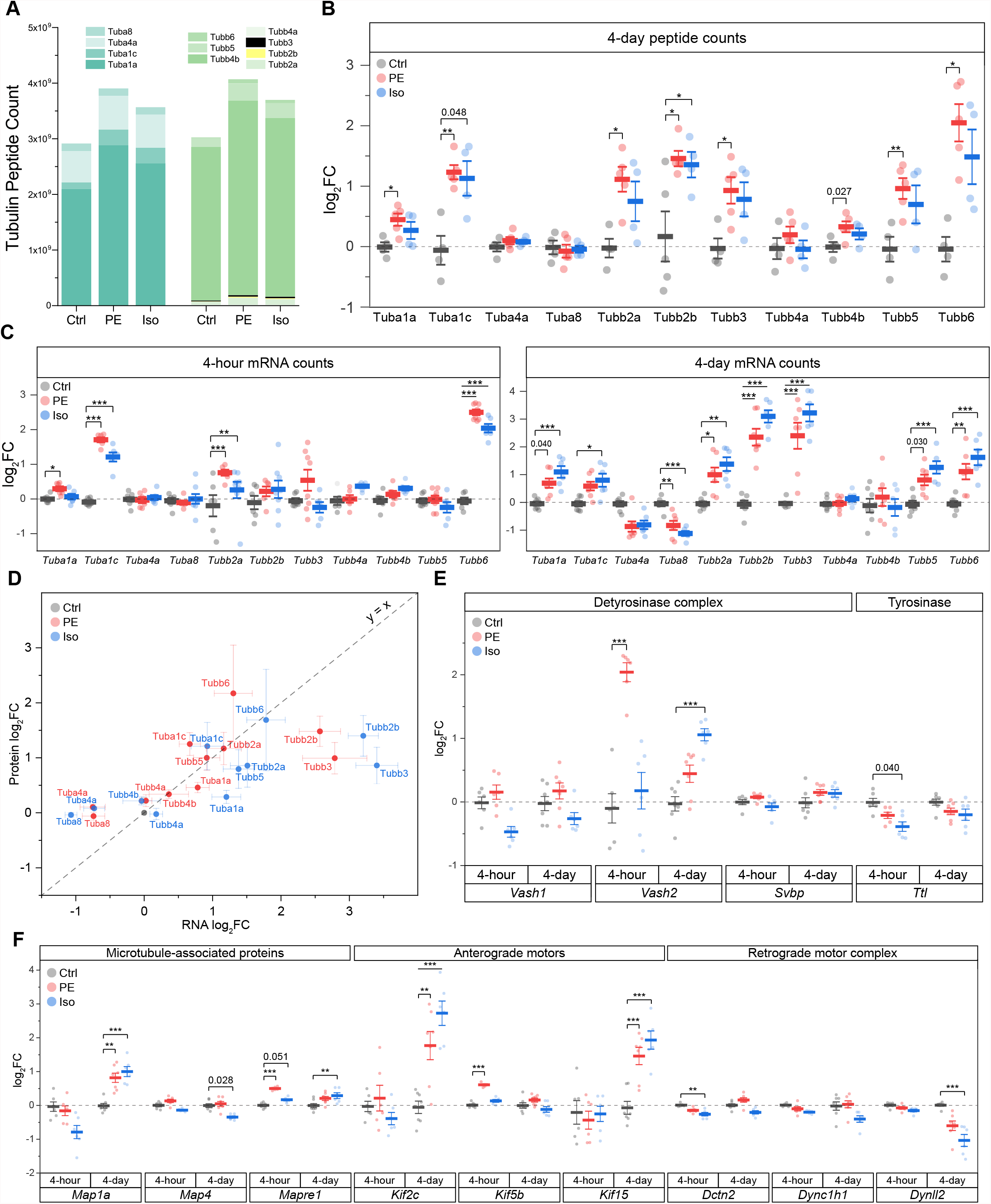
Differential expression of tubulin isoforms, modifying enzymes and MAPs during the onset and establishment of hypertrophy. **(A)** MS counts of unique peptides of detectable αβ-tubulin isoforms at 4-day. For all following box plots, whiskers represent ± 1SEM and bolded line represents mean. **(B)** Relative log_2_fold-change of αβ-tubulin isoforms peptide counts at 4-day (n = Ctrl:5, PE:5, Iso:4); * represents p-value from Welch-corrected two-tailed two-sample t-test on non-log data < 0.025 (Bonferroni-corrected for two comparisons), ** represents p < 0.01, and *** represents p < 0.001. **(C)** relative log_2_fold-change of nCounter mRNA counts of detectable tubulin isoforms at 4-hour **(left)** (n=6) and 4-day **(right)** (n = Ctrl:7, PE:7, Iso:6); * represents Bonferroni adjusted (for 50 genes) p-value < 0.025, ** represents adj_p < 0.01, and *** represents adj_p < 0.001 (see Methods & Materials for more statistical details). **(D)** Scatter-plot of log_2_fold-change of mRNA on x-axis and log_2_fold-change of protein on y-axis at 4-day; whiskers represent ± 1SEM and y = x represents a proportionate change between mRNA and peptide. Relative log_2_fold-change of nCounter mRNA counts of **(E)** detyrosinase complex and tyrosinase, & **(F)** MAPs, anterograde, & retrograde motors at 4-hour (n=6) and 4-day (n = Ctrl:7, PE:7, Iso:6); * represents Bonferroni adjusted (for 50 genes) p-value < 0.025, ** represents adj_p < 0.01, and *** represents adj_p < 0.001 (see Methods for more statistical details).

### Transcriptional analysis of αβ-tubulin isoforms, tubulin modifying enzymes, and MAPs during the induction and establishment of hypertrophy

We next examined the contribution of transcriptional changes to the protein and network level microtubule remodeling at 4-hour and 4-day. To this end, we utilized NanoString analysis of total RNA using another set of 47 genes that includes tubulin isoforms, tubulin modifying enzymes, and MAPs.

While tubulin protein content was unchanged 4-hour after adrenergic stimulation, we noted significant upregulation of several tubulin transcripts with both PE and Iso treatment at this stage, including *Tuba1c, Tubb2a and Tubb6*, with additional and more robust upregulation of *Tubb2b and Tubb3* by day 4 (Fig. 5C). Consistent with proteomics assessments, *Tuba4a* and *Tuba8* were either unchanged or even downregulated upon PE and Iso treatment.

Regardless of the directionality of response, specific tubulin isoforms generally responded similarly to either adrenergic stimulus (Fig. 5C). Further, in contrast to what was observed in advanced HF, transcript levels were also well-correlated with protein abundance across most isoforms at the 4-day time point (R^2^ = 0.38, slope = 0.20, p = 1.4e-4) (Fig. 5D). Consistent with protein expression lagging transcriptional regulation, the four isoforms (Tuba4a, Tuba8, Tubb2b, Tubb3) that displayed the greatest deviation in the change in the mRNA relative to the change in the protein levels (i.e. located furthest away from the y = x line when plotting log_2_FC in mRNA vs protein levels) were transcripts that showed delayed regulation; these isoforms were unchanged after 4-hour but differentially expressed by 4-day. Consistent upregulation at the transcript and protein level was seen for Tuba1c, Tubb2a, Tubb2b, Tubb3, and Tubb6. Combined with the early upregulation of tubulin transcripts, this data indicates that increased tubulin mRNA at least partly underlies the isoform-specific increase in tubulin protein, and therefore tubulin mass, that is necessary for hypertrophic remodeling(Sato et al. 1997; Tsutsui et al. 1999; Scarborough et al. 2021).

We noted several additional transcriptional changes of tubulin modifying enzymes and MAPs that may bear relevance to cardiac remodeling and warrant further investigation (Fig. 5E-F; S. Fig. 4). These include: (1) *Vash2*, which encodes a tubulin detyrosinase, exhibited the greatest differential expression among the 47 assessed transcripts at the 4-hr PE time point; this may contribute to the robust early induction of detyrosination in this group (2) Early upregulation of *Kif5b* in PE(Tigchelaar et al. 2016) after 4-hour, which encodes the primary transport kinesin heavy chain 1 implicated in mRNA transport during myocyte growth(Scarborough et al. 2021); (3) upregulation of *Mapre1* in both PE and Iso at 4-hour, which encodes a member of microtubule associated protein RP/EB family of +TIP tracking protein that guides microtubule growth; (4) Robust upregulation of *Kif15* in both PE and Iso by day 4, which encodes a kinesin family member implicated in stabilizing parallel growing microtubules; (5) induction of *Map1a* in both PE and Iso at 4-day, which encodes a stabilizing structural MAP.

All tubulin-associated transcript volcano plots (S. Fig. 4) were asymmetric, tending to show a greater degree of upregulated than downregulated genes, implying a generalized induction of a tubulin-associated program at 4-day. This was particularly evident in the PE groups, and with progressive upregulation from the 4-hour to 4-day time point. There were, however, notable down-regulated transcripts. While kinesin isoforms, which encode plus-end directed anterograde motors, were generally upregulated in treated groups, transcripts encoding subunits of the dynein/dynactin minus-end directed motor (*Dynll2, Dync1h1, Dctn2)* were either downregulated or unchanged (Fig. 5F). This preferential induction of anterograde motors would bias trafficking toward the microtubule plus-end and away from the minus-end, which has implications for directed cardiac growth and for autophagic flux, which requires minus-end directed transport (McLendon et al. 2014). We also noted the early downregulation of enzymes involved in the polyglutamylation cycle, such as cytosolic carboxypeptidase 5 (*Ccp5*) and TTL-like family members 1 and 5 (*Ttll1/5*), which were all reduced in PE and Iso at the 4-hour time point (S. Fig. 4).

To determine the conservation of these tubulin-associated transcriptional responses across varied hypertrophic stimuli, we compared our data with publicly available RNA sequencing datasets from two separate studies that examined early time-points following pressure-overload and angiotensin II induced hypertrophy. While data is not available for all transcripts, transcripts reported across studies demonstrate well-conserved transcriptional signatures at both early (hours) and later (days) timepoints (S. Fig. 5), including the consistent upregulation of most αβ-tubulin isoforms but with the notable downregulation of *Tuba4a* and *Tuba8*.

### Transcriptional and autoregulatory mechanisms underlie isoform-specific increases in αβ-tubulin mRNA

The above transcriptional and proteomic profiling indicates that the upregulation of tubulin mRNAs is an early driver of microtubule proliferation during the development of hypertrophy. This may arise from two non-exclusive mechanisms – (1) increased transcription or (2) decreased autoregulation (i.e., autoactivation). To this end, we utilized the tubulin isoform and location -specific approach outlined above to interrogate the mechanism of tubulin upregulation during cardiac hypertrophy. Overall, we observed in almost all our tested tubulin isoforms exonic level increases that exceed intronic level increases, suggesting a generalized autoactivation of tubulin isoforms driven by microtubule stabilization (Fig. 6). The most prominent cases of autoactivation are that of *Tubb2b*, whose increase in transcript level is solely through an increase in exonic level at both 4-hour and 4-day, and *Tuba1a*, whose immediate response at 4-hour was through an increase in exonic level with no change in intronic level (Fig. 6A). Additionally, in a subset of the tubulin isoforms – *Tuba1b, Tubb2a, Tubb5*, and *Tubb6 –* we observed robust increases in the intronic levels that could be explained by the direct transcriptional activation of the hypertrophic stimuli (Fig. 6B, S. Fig. 6). Interestingly, despite a generalized upregulation and autoactivation of tubulin isoforms in the early stages of hypertrophy, *Tuba4a* and *Tuba8* are downregulated and autoinhibited, respectively. These data collectively show that αβ-tubulin mRNA is controlled in an isoform-specific and time-dependent fashion through both transcriptional and autoregulatory mechanisms to rewrite the tubulin code during cardiac remodeling.

**Figure 6.**
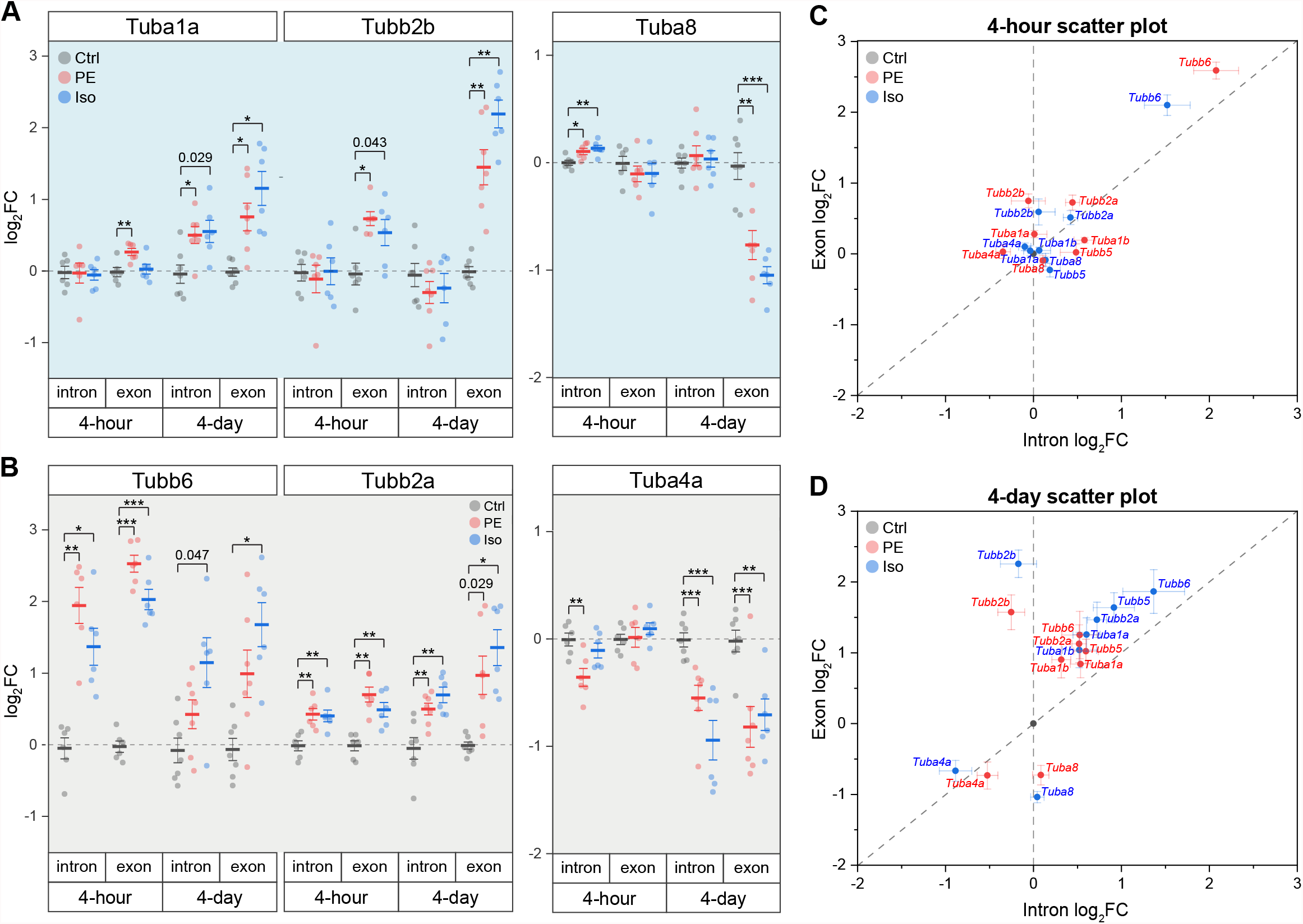
Tubulin isoforms are differentially regulated at the mRNA level through isoform-specific transcription and/or autoregulation during cardiac hypertrophy. Relative log_2_fold-change of mRNA counts of αβ-tubulin isoforms that are predominantly regulated through **(A)** autoregulation, or **(B)** transcription (n = 4h: 6, 4d: Ctrl:7, PE:7, Iso:6); * represents p-value from Welch-corrected two-tailed two-sample t-test on non-log data < 0.025 (Bonferroni-corrected for two comparisons), ** represents p < 0.01, and *** represents p < 0.001. Scatter-plots of relative log_2_fold-change of intron on x-axis and exon on y-axis at **(C)** 4-hour and **(D)** 4-day; whiskers represent ± 1SEM.

## Discussion

In this work we combined transcriptomic and proteomic assessments in advanced heart failure samples and temporally well-defined murine models of cardiac remodeling to understand how a dense and modified microtubule network is achieved. Among other observations expanded upon below, we arrive at three primary conclusions: 1) tubulin autoregulation is operant in the heart and represses tubulin isoform mRNA expression in HF, contributing to the observed discrepancy between RNA and protein levels in HF; 2) the microtubule network is rapidly post-translationally detyrosinated within 4 hours of a hypertrophic stimuli; 3) concomitantly, the abundance of tubulin mRNA is rapidly altered in an isoform-specific fashion through both transcriptional and autoregulatory mechanisms; 4) the time-dependent upregulation of discrete αβ-tubulin transcripts drives an increase in microtubule mass during cardiac hypertrophy.

Combining our work with past literature, we arrive at a sequential model for the formation of a proliferated and stabilized microtubule network in the remodeled heart (S. Fig. 7). Within hours of a hypertrophic stimuli and prior to detectable growth, the microtubule network is detyrosinated (Fig. 3E). Our data indicate that this increase in detyrosination is due to transcriptional (Fig. 5E) or post-translational(Yu et al. 2021) upregulation of the recently identified detyrosinating enzyme complex, as opposed to previously proposed mechanisms such as increased Tuba4a expression (Fig. 5C), increased polymerized or long-lasting microtubule substrate (Fig. 3C-D), or decreased TTL expression (S. Fig. 3B, E). Detyrosination serves as a network stabilizer to protect microtubules from breaking down by regulating their interaction with effector proteins(Peris et al. 2009; Chen et al. 2021). Microtubule stabilization, in turn, shuttles free tubulin into the polymerized microtubule pool, triggering autoactivation that increases tubulin mRNA stability and translation. How autoregulation may achieve isoform-specificity is not understood, although indicated by our data (see *Tubb2b* vs. *Tuba8*, Fig. 6F). In concert with post-transcriptional upregulation of tubulin mRNA, increased transcription of several isoforms concomitantly increases tubulin mRNA. Independent of the mode of upregulation, tubulin mRNAs appear to be efficiently translated, as mRNA levels are well correlated with peptide abundance across tubulin isoforms (Fig. 5D). As the stimuli persists and the heart enlarges, the newly translated tubulin is integrated into the microtubule network, resulting in increased microtubule mass and additional substrate for post-translational modifications (Fig. 4).

Insights into tubulin isoforms in muscle biology have pointed towards the potential detrimental effects of specific isoforms in muscle pathologies; for example, TUBB6 is upregulated in dystrophic skeletal muscles, and it contributes to microtubule disorganization and altered muscle regeneration in muscular dystrophy(Randazzo et al. 2019), and elevated TUBA4A in human cardiomyopathy contributes to the increased detyrosination that impedes myocyte function(Chen et al. 2018; Schuldt et al. 2020) (Fig. 5B). Strikingly, when we examine publicly available transcriptomic and proteomic data from chronically hypertrophied or failing human hearts (Fig. 1A), we observe an inverse relationship between the transcript and the protein levels of all αβ-tubulin isoforms. It is worth noting that TUBA8 behaves as an outlier, the lone tubulin transcript that is consistently *increased* in HF while the protein level is consistently *decreased*. Intriguingly, *Tuba8* was also the sole isoform to clearly escape autoactivation (and appear seemingly autoinhibited) during early hypertrophic remodeling (Fig. 6F). We have no current explanation for how or why Tuba8 shows unique regulation in both settings. In contrast to inverse relationship in HF, we observed that during the establishment of hypertrophy, transcript and protein levels are highly correlated, suggesting an uncoupling of transcript and protein levels that occurs later in the course of cardiac remodeling. Chronic, robust microtubule stabilization and increased tubulin lifetime could account for the stably elevated tubulin protein content despite persistent autoinhibition that we observe in HF.

Our analysis permits the temporal evaluation of several cytoskeletal- or hypertrophy-associated factors at distinct stages representing the onset and establishment of cardiac hypertrophy. Beyond the key conclusions listed above, several additional observations on cytoskeletal remodeling are of note. The association of the microtubule network with motor proteins such as kinesins alters its mechano-biochemical properties as well as its density. As an example, Kif15 (kinesin-12) has been shown to cross-link nearby parallel microtubules, causing them to bundle, and subsequently decreases the catastrophic events of dynamic microtubules(Drechsler and McAinsh 2016). Interestingly, during both PE and Iso -induced hypertrophy, *Kif15* is upregulated, suggesting that *Kif15* could contribute to microtubule network densification. Kif5b (Kinesin-1), the predominant anterograde motor in the heart, was previously reported to be increased in PE induced-hypertrophy of neonatal rat ventricular cardiomyocytes(Tigchelaar et al. 2016). We observed similar and rapid increase in Kif5b transcript and protein levels in our hypertrophy models (Fig. 5F, S. Fig. 3C, F). Kinesin-1 was recently identified to be required for the distribution of mRNA and ribosomes that enables cardiomyocyte hypertrophy (Scarborough et al. 2021), and past work indicates that kinesin-1 prefers to transport cargo along detyrosinated microtubule tracks(Kaul et al. 2014). Meanwhile, the dynein/dynactin retrograde motor protein complex (transcriptionally downregulated, Fig. 5F), prefers tyrosinated microtubule tracks(Nirschl et al. 2016). Taking together, these observations suggest that the heart both rapidly induces its primary anterograde transport motor and remodels its preferred tracks in response to a hypertrophic stimulus.

Our findings indicate that rapid transcriptional, autoregulatory, and post-translational mechanisms remodel the microtubule network following a hypertrophic stimulus. Contextualized with past literature, these changes will support the ability of the microtubule network to bear increased mechanical load, facilitate mechanotransduction, and enhance transport of the translational machinery that is required for growth. In summary, the data points towards a concerted and adaptive response to establish hypertrophy, and we provide a resource for further investigation into the diverse roles of microtubules in cardiac remodeling.

## Abbreviations and Acronyms

HF: Heart Failure
PTM: Post-Translational Modification
MAP: Microtubule-Associated Proteins
αTAT1: α-Tubulin acetyltransferase 1
VASH1/2: vasohibins 1 & 2
HW: Heart Weight
TL: Tibia Length
LV: Left-Ventricle
Ctrl: Vehicle control
PE: Phenylephrine
Iso: Isoproterenol
IEG: Immediate Early Gene
DEG: Differentially Expressed Gene

## Conflict of Interest

The authors declare no conflict of interest.

## Author Contributions

SP and BP designed the study. SP, KU, CC, MC, JG and KB performed data acquisition and analysis. SP and BP wrote the manuscript, and all authors assisted in editing.

## Funding

Funding for this work came was provided by the National Institute of Health (NIH) R01s-HL133080 and HL149891 to B. Prosser, T32 HL007843 to K. Uchida, by the Fondation Leducq Research Grant no. 20CVD01 to B. Prosser, and by the Center for Engineering Mechanobiology to B. Prosser through a grant from the National Science Foundation’s Science and Technology program: 15-48571.

## Acknowledgment

We thank the Mouse Cardiovascular Phenotyping Core of the Cardiovascular Institute at the University of Pennsylvania for the echocardiography, the Quantitative Proteomics Resource Core of School of Medicine at the University of Pennsylvania for the mass spectrometry, the Penn Center for Musculoskeletal Disorders Histology Core (P30-AR069619) & the Genomics Core at the Wistar Institute for the NanoString analyses. Some of the figures were created by BioRender.com.

## Notes

### Competing Interest Statement

The authors have declared no competing interest.

## References

Aillaud C, Bosc C, Peris L, Bosson A, Heemeryck P, Dijk JV, et al. Vasohibins/SVBP are tubulin carboxypeptidases (TCPs) that regulate neuron differentiation. Science. 2017;358(6369):1448–53.

Aillaud C, Bosc C, Saoudi Y, Denarier E, Peris L, Sago L, et al. Evidence for new C-terminally truncated variants of α- and β-tubulins. Mol Biol Cell. 2016;27(4):640–53.

Baehr A, Umansky KB, Bassat E, Jurisch V, Klett K, Bozoglu T, et al. Agrin Promotes Coordinated Therapeutic Processes Leading to Improved Cardiac Repair in Pigs. Circulation. 2020;142(9):868–81.

Barton PJR, Birks EJ, Felkin LE, Cullen ME, Koban MU, Yacoub MH. Increased expression of extracellular matrix regulators TIMP1 and MMP1 in deteriorating heart failure. J Hear Lung Transplant. 2003;22(7):738–44.

Bassat E, Mutlak YE, Genzelinakh A, Shadrin IY, Umansky KB, Yifa O, et al. The extracellular matrix protein agrin promotes heart regeneration in mice. Nature. 2017;547(7662):179–84.

Caporizzo MA, Chen CY, Bedi K, Margulies KB, Prosser BL. Microtubules Increase Diastolic Stiffness in Failing Human Cardiomyocytes and Myocardium. Circulation. 2020;141(11):902–15.

Caporizzo MA, Chen CY, Prosser BL. Cardiac microtubules in health and heart disease. Exp Biol Med. 2019;244(15):1255–72.

Caporizzo MA, Chen CY, Salomon AK, Margulies KB, Prosser BL. Microtubules Provide a Viscoelastic Resistance to Myocyte Motion. Biophys J. 2018;115(9):1796–807.

Chen CY, Caporizzo MA, Bedi K, Vite A, Bogush AI, Robison P, et al. Suppression of detyrosinated microtubules improves cardiomyocyte function in human heart failure. Nat Med. 2018;24(8):1225–33.

Chen J, Kholina E, Szyk A, Fedorov VA, Kovalenko I, Gudimchuk N, et al. α-tubulin tail modifications regulate microtubule stability through selective effector recruitment, not changes in intrinsic polymer dynamics. Dev Cell. 2021;

Cheng G, Takahashi M, Shunmugavel A, Wallenborn JG, DePaoli-Roach AA, Gergs U, et al. Basis for MAP4 Dephosphorylation-related Microtubule Network Densification in Pressure Overload Cardiac Hypertrophy. J Biol Chem. 2010;285(49):38125–40.

Cheng G, Zile MR, Takahashi M, Baicu CF, Bonnema DD, Cabral F, et al. A direct test of the hypothesis that increased microtubule network density contributes to contractile dysfunction of the hypertrophied heart. Am J Physiol-heart C. 2008;294(5):H2231–41.

Drechsler H, McAinsh AD. Kinesin-12 motors cooperate to suppress microtubule catastrophes and drive the formation of parallel microtubule bundles. Proc National Acad Sci. 2016;113(12):E1635–44.

Fassett J, Xu X, Kwak D, Zhu G, Fassett EK, Zhang P, et al. Adenosine kinase attenuates cardiomyocyte microtubule stabilization and protects against pressure overload-induced hypertrophy and LV dysfunction. J Mol Cell Cardiol. 2019;130:49–58.

Fassett JT, Xu X, Hu X, Zhu G, French J, Chen Y, et al. Adenosine regulation of microtubule dynamics in cardiac hypertrophy. Am J Physiol-heart C. 2009;297(2):H523–32.

Friedrich FW, Wilding BR, Reischmann S, Crocini C, Lang P, Charron P, et al. Evidence for FHL1 as a novel disease gene for isolated hypertrophic cardiomyopathy. Hum Mol Genet. 2012;21(14):3237–54.

Gasic I, Boswell SA, Mitchison TJ. Tubulin mRNA stability is sensitive to change in microtubule dynamics caused by multiple physiological and toxic cues. Plos Biol. 2019;17(4):e3000225.

Gasic I, Mitchison TJ. Autoregulation and repair in microtubule homeostasis. Curr Opin Cell Biol. 2019;56:80–7.

Hayata N, Fujio Y, Yamamoto Y, Iwakura T, Obana M, Takai M, et al. Connective tissue growth factor induces cardiac hypertrophy through Akt signaling. Biochem Bioph Res Co. 2008;370(2):274–8.

Kalebic N, Sorrentino S, Perlas E, Bolasco G, Martinez C, Heppenstall PA. αTAT1 is the major α-tubulin acetyltransferase in mice. Nat Commun. 2013;4(1):1962.

Kaul N, Soppina V, Verhey KJ. Effects of α-Tubulin K40 Acetylation and Detyrosination on Kinesin-1 Motility in a Purified System. Biophys J. 2014;106(12):2636–43.

Li L, Zhang Q, Zhang X, Zhang J, Wang X, Ren J, et al. Microtubule associated protein 4 phosphorylation leads to pathological cardiac remodeling in mice. Ebiomedicine. 2018;37:221–35.

McLendon PM, Ferguson BS, Osinska H, Bhuiyan MdS, James J, McKinsey TA, et al. Tubulin hyperacetylation is adaptive in cardiac proteotoxicity by promoting autophagy. Proc National Acad Sci. 2014;111(48):E5178–86.

Nieuwenhuis J, Adamopoulos A, Bleijerveld OB, Mazouzi A, Stickel E, Celie P, et al. Vasohibins encode tubulin detyrosinating activity. Science. 2017;358(6369):1453–6.

Nirschl JJ, Magiera MM, Lazarus JE, Janke C, Holzbaur ELF. α-Tubulin Tyrosination and CLIP-170 Phosphorylation Regulate the Initiation of Dynein-Driven Transport in Neurons. Cell Reports. 2016;14(11):2637–52.

Paturle-Lafanechère L, Manier M, Trigault N, Pirollet F, Mazarguil H, Job D. Accumulation of delta 2-tubulin, a major tubulin variant that cannot be tyrosinated, in neuronal tissues and in stable microtubule assemblies. J Cell Sci. 1994;107 (Pt 6):1529–43.

Peris L, Wagenbach M, Lafanechère L, Brocard J, Moore AT, Kozielski F, et al. Motor-dependent microtubule disassembly driven by tubulin tyrosination. J Cell Biol. 2009;185(7):1159–66.

Portran D, Schaedel L, Xu Z, Théry M, Nachury MV. Tubulin acetylation protects long-lived microtubules against mechanical ageing. Nat Cell Biol. 2017;19(4):391–8.

Prosser BL, Ward CW, Lederer WJ. X-ROS Signaling: Rapid Mechano-Chemo Transduction in Heart. Science. 2011;333(6048):1440–5.

Randazzo D, Khalique U, Belanto JJ, Kenea A, Talsness DM, Olthoff JT, et al. Persistent upregulation of the β-tubulin tubb6, linked to muscle regeneration, is a source of microtubule disorganization in dystrophic muscle. Hum Mol Genet. 2019;28(7):1117–35.

Sato H, Nagai T, Kuppuswamy D, Narishige T, Koide M, Menick DR, et al. Microtubule Stabilization in Pressure Overload Cardiac Hypertrophy. J Cell Biol. 1997;139(4):963–73.

Scarborough EA, Uchida K, Vogel M, Erlitzki N, Iyer M, Phyo SA, et al. Microtubules orchestrate local translation to enable cardiac growth. Nat Commun. 2021;12(1):1547.

Schuldt M, Pei J, Harakalova M, Dorsch LM, Schlossarek S, Mokry M, et al. Proteomic and Functional Studies Reveal Detyrosinated Tubulin as Treatment Target in Sarcomere Mutation-Induced Hypertrophic Cardiomyopathy. Circulation Hear Fail. 2020;14(1):e007022.

Steffensen LB, Rasmussen LM. A role for collagen type IV in cardiovascular disease? Am J Physiol-heart C. 2018;315(3):H610–25.

Swiatlowska P, Sanchez-Alonso JL, Mansfield C, Scaini D, Korchev Y, Novak P, et al. Short-term angiotensin II treatment regulates cardiac nanomechanics via microtubule modifications. Nanoscale. 2020;12(30):16315–29.

Tigchelaar W, Jong AM de, Bloks VW, Gilst WH van, Boer RA de, Silljé HHW. Hypertrophy induced KIF5B controls mitochondrial localization and function in neonatal rat cardiomyocytes. J Mol Cell Cardiol. 2016;97:70–81.

Tsutsui H, Ishihara K, Cooper G. Cytoskeletal role in the contractile dysfunction of hypertrophied myocardium. Science. 1993;260(5108):682–7.

Tsutsui H, Tagawa H, Cooper G, Takeshita A. Role of Microtubules in the Transition from Compensated Cardiac Hypertrophy to Failure. Heart Fail Rev. 1999;4(4):311–8.

Vistnes M, Aronsen JM, Lunde IG, Sjaastad I, Carlson CR, Christensen G. Pentosan Polysulfate Decreases Myocardial Expression of the Extracellular Matrix Enzyme ADAMTS4 and Improves Cardiac Function In Vivo in Rats Subjected to Pressure Overload by Aortic Banding. Plos One. 2014;9(3):e89621.

Xu Z, Schaedel L, Portran D, Aguilar A, Gaillard J, Marinkovich MP, et al. Microtubules acquire resistance from mechanical breakage through intralumenal acetylation. Science. 2017;356(6335):328–32.

Yu X, Chen X, Amrute-Nayak M, Allgeyer E, Zhao A, Chenoweth H, et al. MARK4 controls ischaemic heart failure through microtubule detyrosination. Nature. 2021;1–6.

[dataset] Liu Y, Morley M, Brandimarto J, Hannenhalli S, Hu Y, Ashley EA, Tang WH, Moravec CS, Margulies KB, Cappola TP, Li M; MAGNet consortium. RNA-Seq identifies novel myocardial gene expression signatures of heart failure. Genomics. 2015 Feb;105(2):83-9. doi: 10.1016/j.ygeno.2014.12.002.

[dataset] Schuldt M, Pei J, Harakalova M, Dorsch LM, Schlossarek S, Mokry M, Knol JC, Pham TV, Schelfhorst T, Piersma SR, Remedios C dos, Dalinghaus M, Michels M, Asselbergs FW, Moutin M-J, Carrier L, Jimenez CR, Velden J van der, Kuster DWD. Proteomic and Functional Studies Reveal Detyrosinated Tubulin as Treatment Target in Sarcomere Mutation-Induced Hypertrophic Cardiomyopathy. Circulation Hear Fail 2020;14:e007022.

[dataset] Doroudgar S, Hofmann C, Boileau E, Malone B, Riechert E, Gorska AA, Jakobi T, Sandmann C, Jürgensen L, Kmietczyk V, Malovrh E, Burghaus J, Rettel M, Stein F, Younesi F, Friedrich UA, Mauz V, Backs J, Kramer G, Katus HA, Dieterich C, Völkers M. Monitoring Cell-Type–Specific Gene Expression Using Ribosome Profiling In Vivo During Cardiac Hemodynamic Stress. Circ Res 2019;125:431–448. doi: 10.1161/CIRCRESAHA.119.314817

[dataset] Bottermann K, Leitner L, Pfeffer M, Nemmer J, Deenen R, Köhrer K, Stegbauer J, Gödecke A. Effect of cardiomyocyte and vascular smooth muscle cell specific KO of p38 MAPKα on gene expression profile of the heart. Gene Expression Omnibus. ID: GSE86074

