## Supplemental Figures for "Transcriptional, post-transcriptional, and post-translational mechanisms rewrite the tubulin code during cardiac hypertrophy and failure"

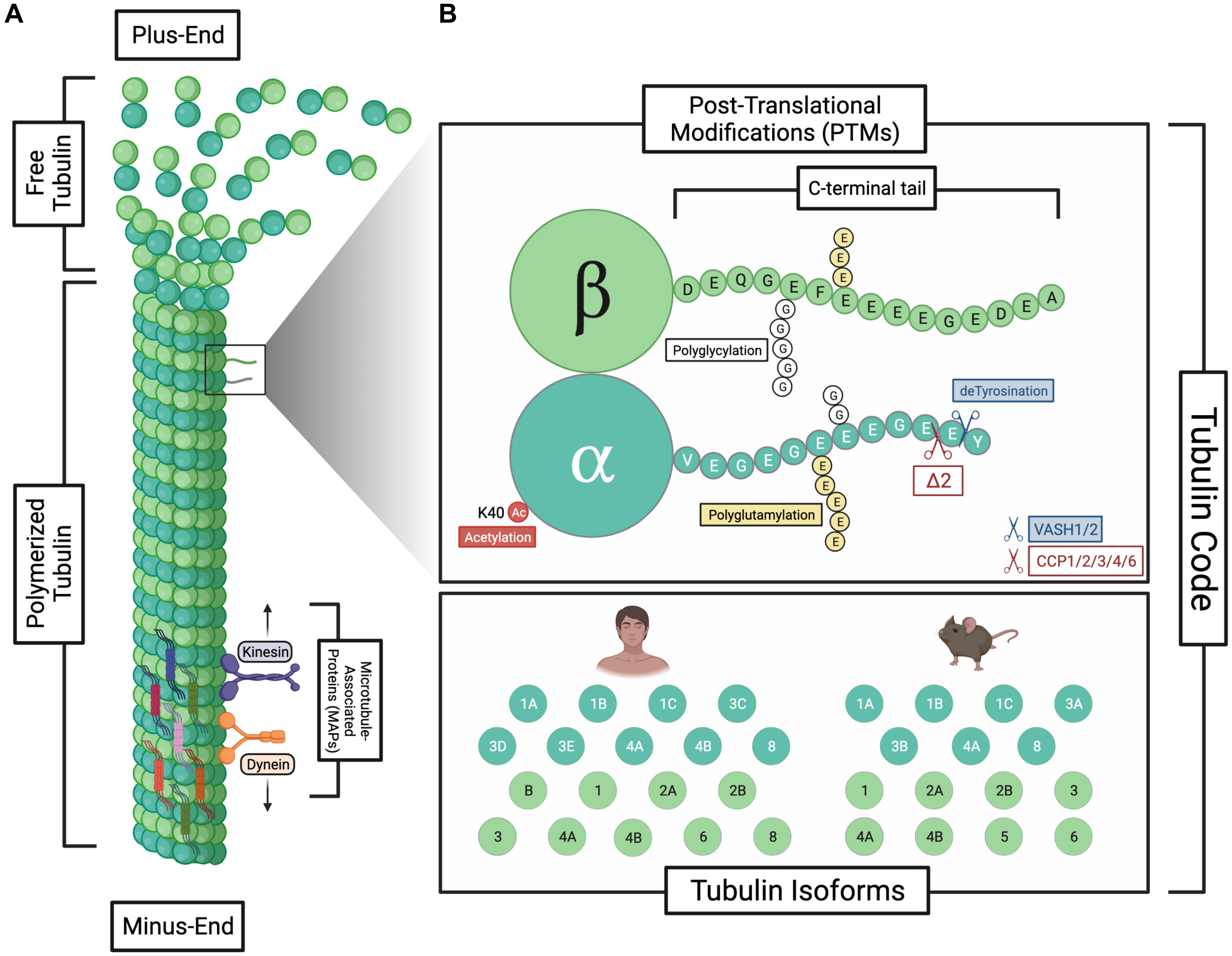


**Supplemental Figure 1.** Summary of the tubulin code. **(A)** Free & polymerized tubulins are in dynamic equilibrium. **(B)** The tubulin code is the permutation of the tubulin post-translational modifications & isoforms.

### Hypertrophy & Fibrosis

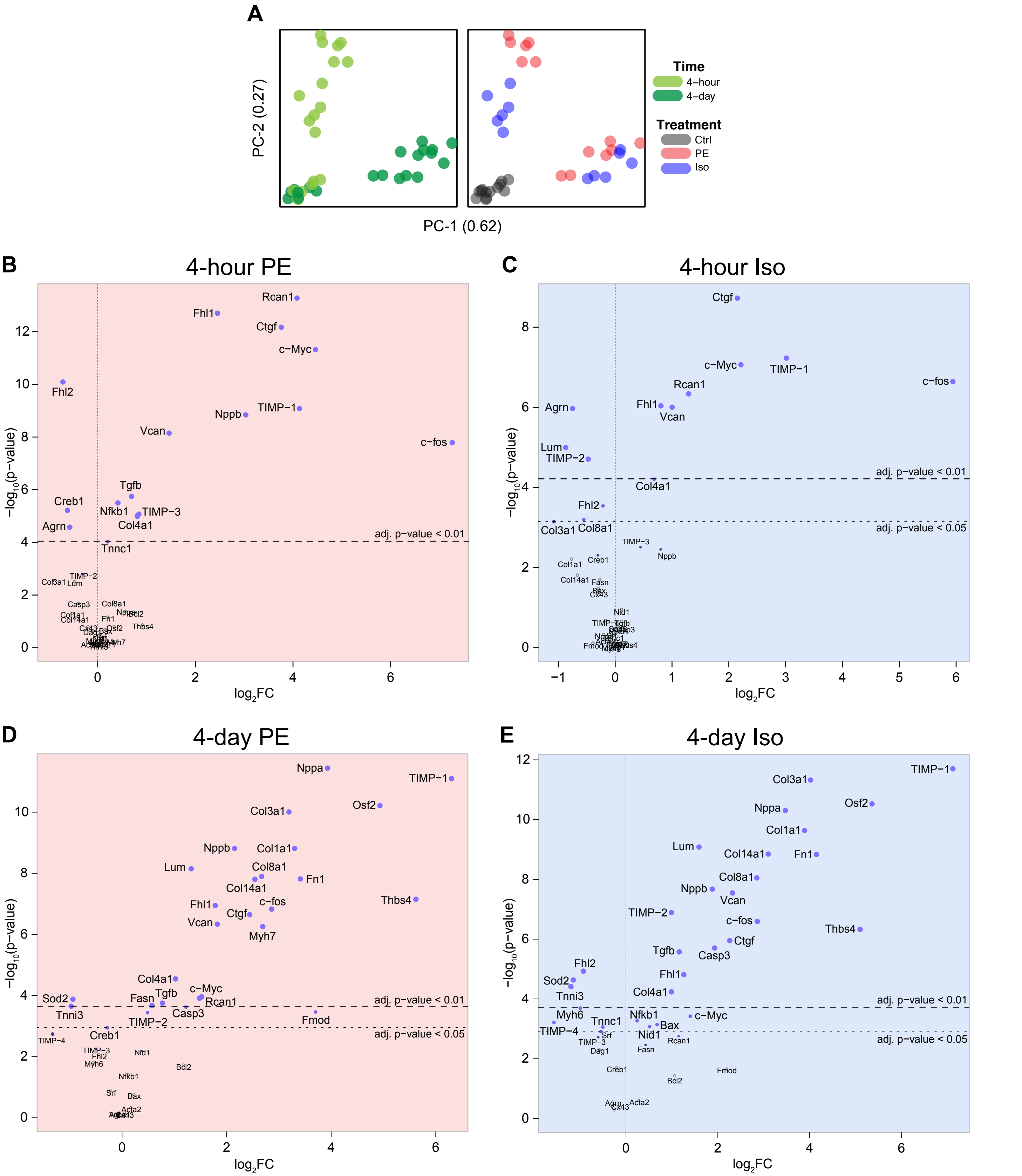


**Supplemental Figure 2. (A)** Principal component analysis (PCA) of nCounter mRNA counts from 4-hour and 4-day mice heart tissues using the hypertrophy & fibrosis panel. Volcano plots of all mRNA counts from the hypertrophy & fibrosis panel for **(B)** 4-hour PE, **(C)** 4-hour Iso, **(D)** 4-day PE, & **(E)** 4-day Iso. (n = 4h:6, 4d: Ctrl:7, PE:7, Iso:6)

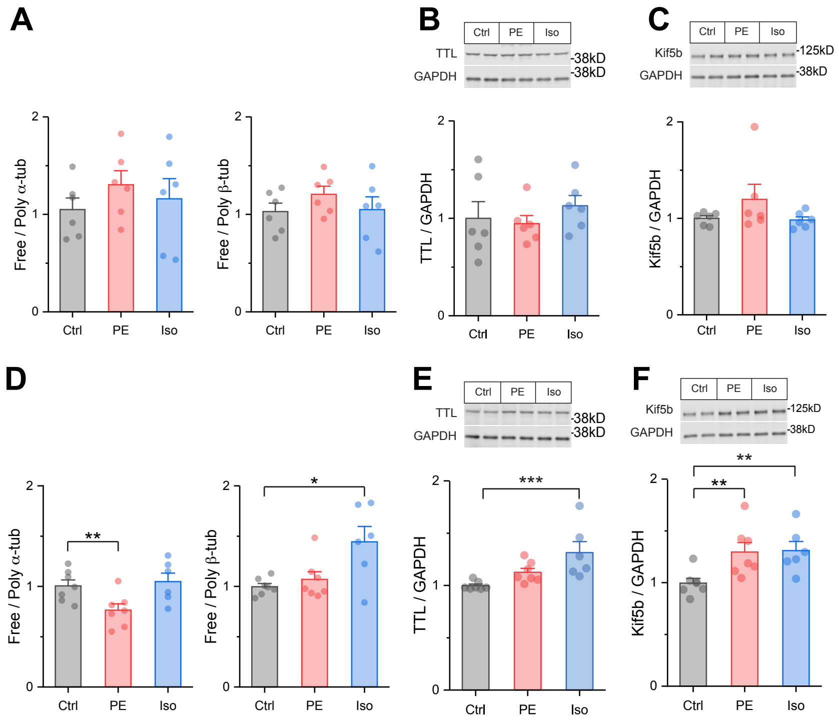


**Supplemental Figure 3.** Free/Poly for **(A, C)** α **(left)** or β **(right)** -tubulins, and **(B, D)** TTL for **(A-B)** 4-hour & **(C-D)** 4-day mice heart tissues (n = 4h:6, 4d: Ctrl:7, PE:7, Iso:6). For all graphs, * represents p-value from Welch-corrected two-tailed two-sample t-test < 0.025, ** represents p < 0.01, and *** represents p < 0.001.

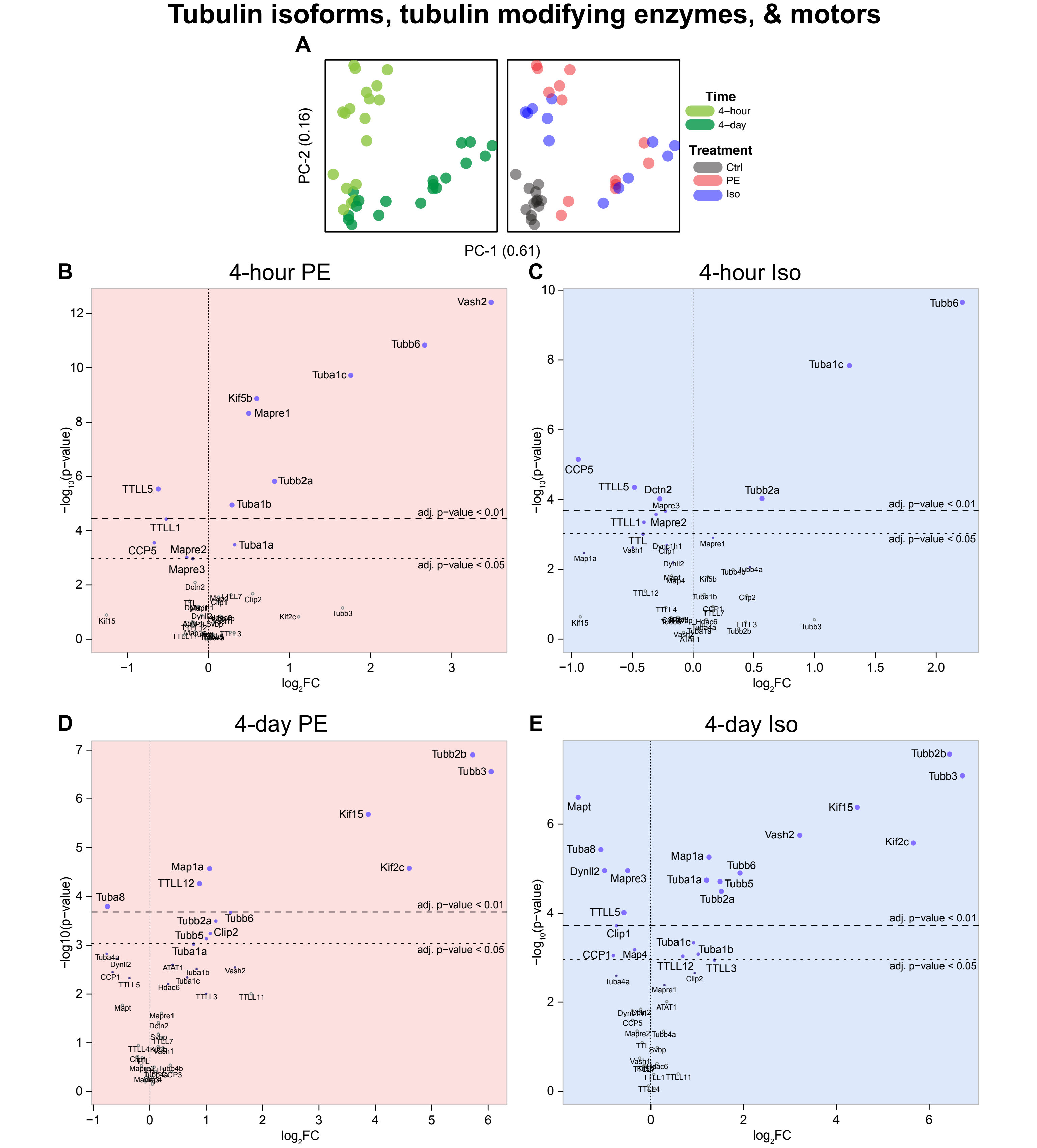


**Supplemental Figure 4. (A)** Principal component analysis (PCA) of nCounter mRNA counts from 4-hour and 4-day mice heart tissues using the tubulin panel. Volcano plots of all mRNA counts from the tubulin panel for **(B)** 4-hour PE, **(C)** 4-hour Iso, **(D)** 4-day PE, & **(E)** 4-day Iso. (n = 4h:6, 4d: Ctrl:7, PE:7, Iso:6)

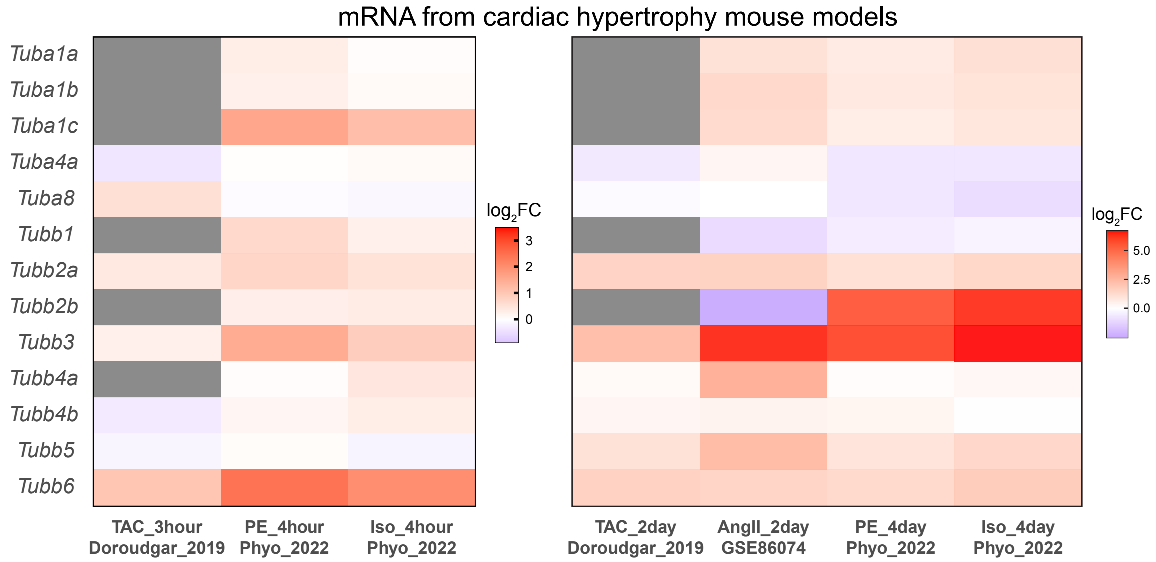


**Supplemental Figure 5.** Previously published & current studies' mRNA data of mouse $\alpha\beta$-tubulin isoforms following different pathological stimulations.

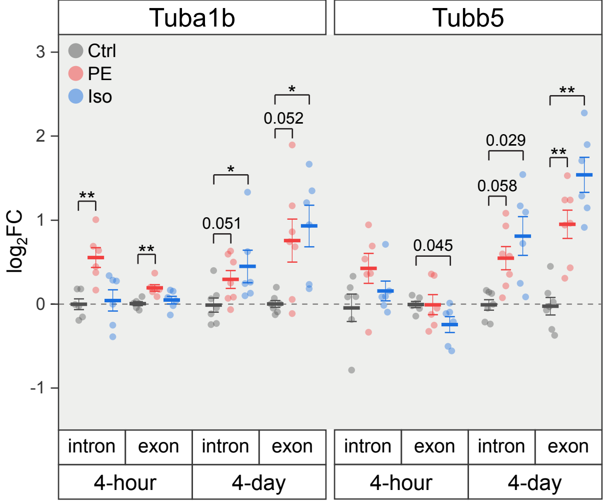


**Supplemental Figure 6.** Unspliced and spliced mRNA counts of Tuba1b and Tubb5 (n = 4h:6, 4d: Ctrl:7, PE:7, Iso:6); * represents p-value from Welch-corrected two-tailed two-sample t-test on non-log data < 0.025 (Bonferroni-corrected for two comparisons), ** represents p < 0.01, and *** represents p < 0.001.

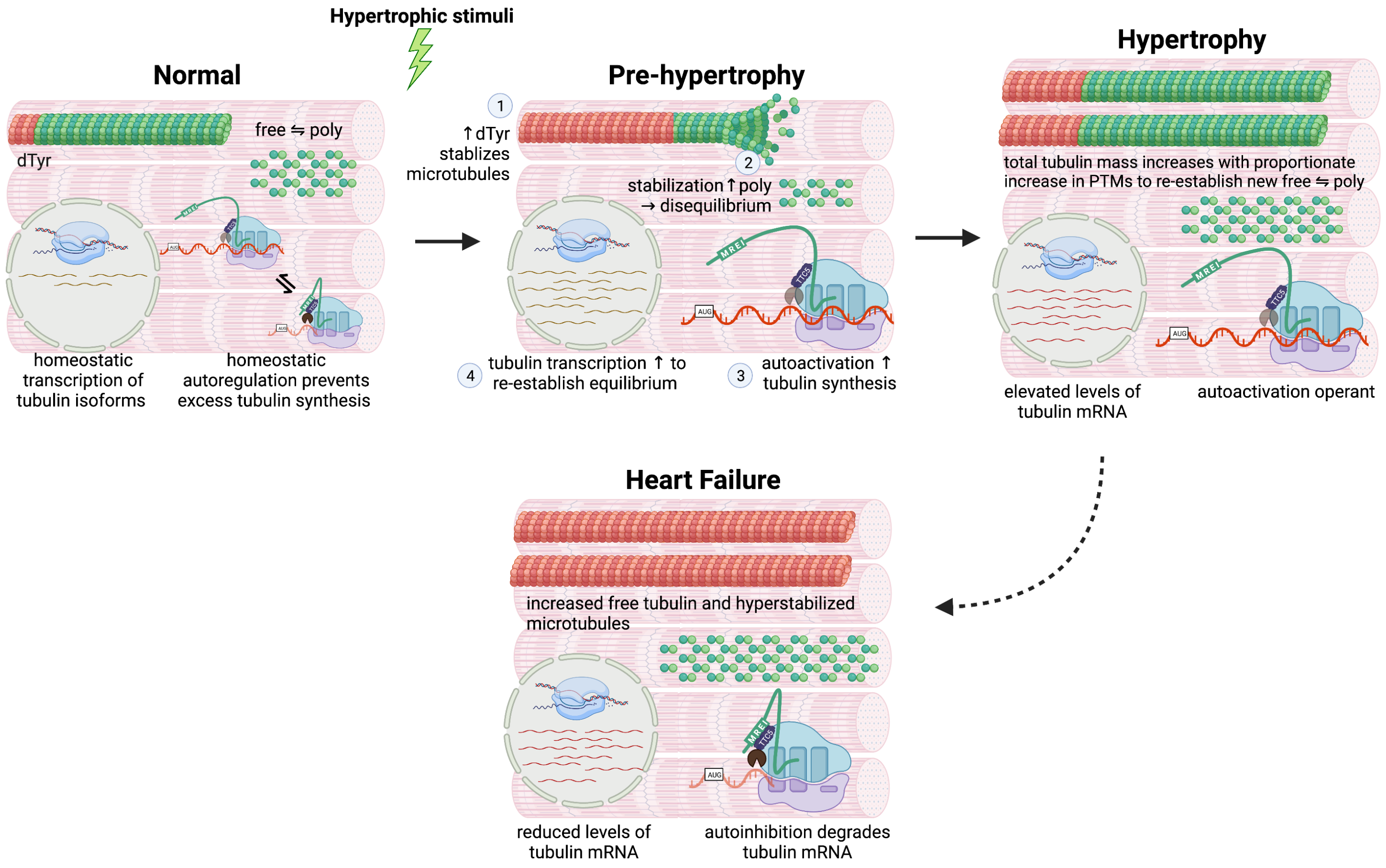


**Supplemental Figure 7.** Schematic model of the summary
